# Neural circuit dynamics of drug-context associative learning in the hippocampus

**DOI:** 10.1101/2021.09.02.458796

**Authors:** Yanjun Sun, Lisa M Giocomo

**Author notes:** Corresponding authors (LMG), (YS).

## Abstract

The environmental context associated with previous drug consumption serves as a potent trigger for relapse to drug use. The mechanism by which existing neural representations of context are modified to incorporate information associated with a given drug however, remains unknown. Using longitudinal calcium imaging in freely behaving mice, we reveal that drug-context associations for psychostimulants and opioids are encoded in a subset of hippocampal neurons. In these neurons, drug context pairing in a conditioned place preference task weakened their spatial coding for the nondrug-paired context, with drug-induced changes to spatial coding predictive of drug-seeking behavior. Furthermore, the dissociative drug ketamine blocked both the drug-induced changes to hippocampal coding and corresponding drug-seeking behavior. Together, this work reveals how drugs of abuse can alter the hippocampal circuit to encode drug-context associations and points to the hippocampus as a key node in the cognitive process of drug addiction and context-induced drug relapse.

## Introduction

A core challenge of long-term recovery from drug addiction is the associated high relapse rate, in which a person returns to drug use after a period of abstinence (*1*). One of the strongest triggers for relapse in both humans and animal models is re-exposure to a drug-associated environmental context (*2–6*). During repeated drug use, a given environmental context is passively associated with the rewarding effects of the drug, such that a previously neutral context may become a conditioned stimulus that can reliably reinstate drug seeking behavior (*7*). In animals, this association can be captured by conditioned place preference (CPP), in which a drug is administered in a specific environmental context, resulting in the animal subsequently spending more time in the drug-paired, compared to the neutral, context (*8, 9*). Converging evidence suggests this type of conditioned learning may rely on multiple memory systems working in parallel, including the nucleus accumbens (NAc), amygdala, and hippocampus (*5, 10–16*) .

In particular, the hippocampus is well positioned to support the encoding of a drug-context association during the conditioned learning (*17–23*). The hippocampus contains place cells that fire in one or few restricted spatial locations in a given environment (*24, 25*). As a population, place cells are proposed to construct a map-like representation for a given environmental context (*25, 26*). Across different spatial environments, or contexts, place cells can show uncorrelated activity, with place fields turning on, off or firing in a new spatial position (*26–37*). These changes in place field activity are collectively referred to as place cell ‘remapping’, which encompasses two phenomena: a change in the firing rate of a place field (rate remapping) and a change in the spatial location of the place field (global remapping) (*38, 39*). The observations of place cell remapping has lent significant support to the theory that the hippocampus contains the neural representations needed to encode the combination of sensory cues and internal variables that define a given spatial context (*38, 40*).

The potential role of the hippocampus in drug-context associative learning is also supported by the sensitivity of place cell representations to the presence of natural reward, such as food or water. Place cells, particularly in the hippocampal subregion CA1, often remap their place fields to cluster near locations associated with rewards, resulting in an overrepresentation of reward locations (*36, 41–47*). Such reward-driven place cell reorganization reflects, at least in part, inputs from the catecholaminergic system (*45*). Recent studies have also reported that cocaine strengthens coupling between hippocampal place cells associated with drug-paired locations and the nucleus accumbens (*18*) and disruption of hippocampal neural representations of a drug-associated context can neutralize drug-induced place preference (*20*). However, the degree to which hippocampal place cell representations maladaptively remap to encode or maintain drug-context associations remains incompletely understood.

Here, we imaged calcium activity in freely moving mice (*48–50*) to examine hippocampal place cell representations over the course of conditioned place preference. We first focused on the effects of methamphetamine (MA), a widely abused psychostimulant that alters catecholaminergic signaling (*51*). We identified a subset of CA1 place cells that remap to represent the drug-paired context after MA conditioning, switching off their activity on the neutral/saline-paired side and sharpening their activity on the MA-paired side of the CPP apparatus. This place cell remapping correlated with the MA-induced place preference behavior and emerged via an experience dependent mechanism. Further, both MA-associated place cell remapping and CPP behavior was blocked with the administration of ketamine, which produces a dissociation-like state in mice (*52*). Finally, we extended these findings to morphine (MO), and report that MO-context associations were encoded through a similar mechanism but recruited a different functionally defined population of place cells. Together, these observations reveal the dynamics of hippocampal place cell remapping in response to drug-context associative learning and provide evidence of strong correspondence between place cell remapping and an animal’s behavioral preference for spatial contexts associated with drug administration.

## Results

### Imaging of CA1 cells in a conditioned place preference (CPP) paradigm

To study the effect of methamphetamine (MA) on the activity of CA1 neurons, we performed in vivo single photon (1P) miniscope calcium imaging in mice during a conditioned place preference paradigm (CPP) (Figure 1). The CPP apparatus consisted of two compartments with distinct colors and visual cues (Figure 1A), which could be connected by a junction in the middle (Figures 1A and 1B). After two days of habituation, mice were allowed access to both compartments of the apparatus for the pre-baseline (pre-bsl) and baseline (bsl) sessions (day 1 = pre-bsl; day 2 = bsl) (Figure 1B). An animal’s natural compartment preference was determined on day 2 (bsl session) (Figure 1B). For conditioning (n = 3 sessions), MA mice were restricted to one of the compartments, with saline injections paired with the naturally preferred side and MA injections paired with the non-preferred side (Figure 1B). One and six days after conditioning, two test sessions (test1 and test2, respectively) were performed to assess post-conditioning place preference (Figure 1B). Control (Ctrl) mice underwent the same protocol but saline injections were paired with both compartments during the conditioning (Figure 1B). In MA mice, we observed significant place preference for the MA-paired compartment in the test sessions (Figure 1C, Methods).

**Figure 1.**
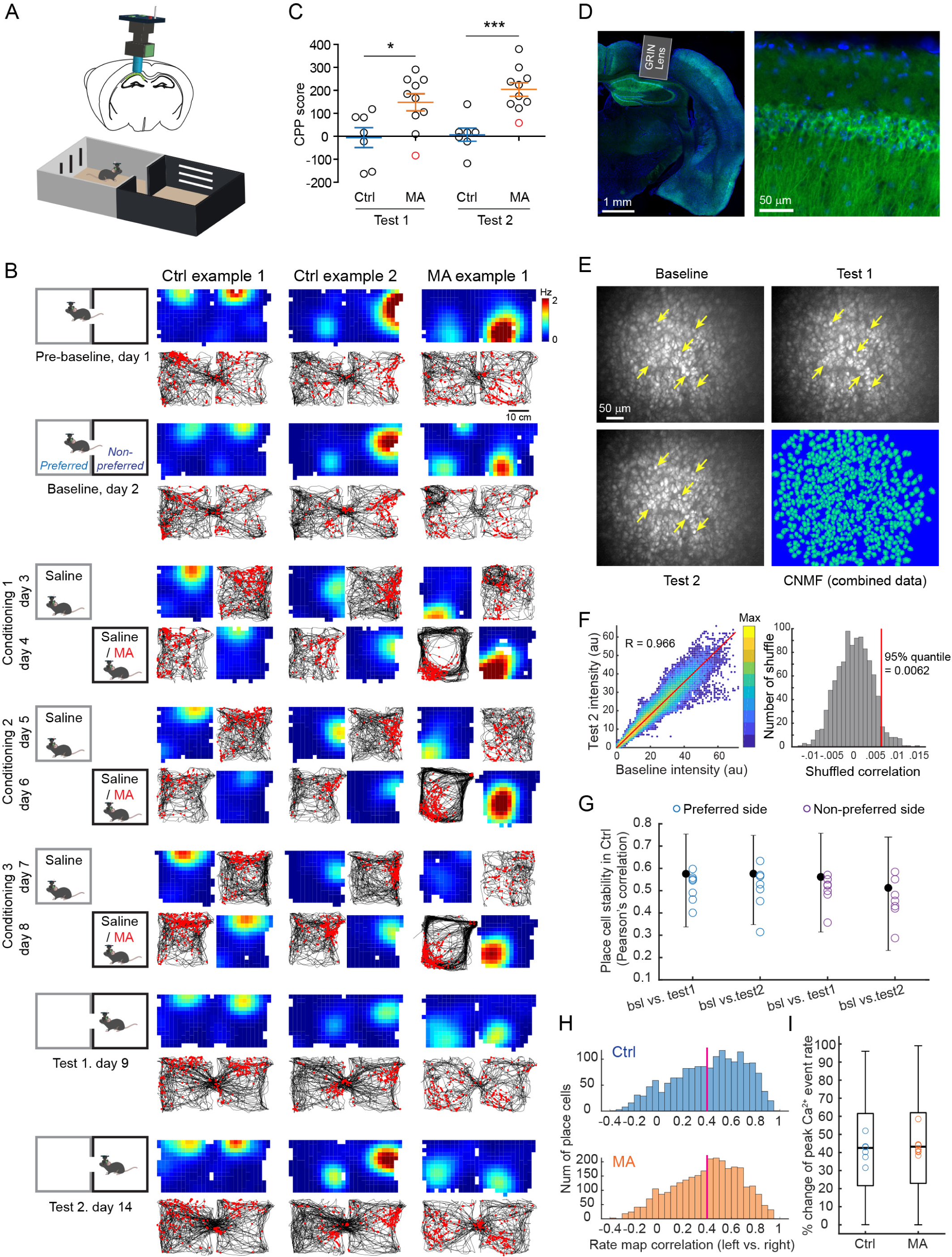
Imaging of CA1 cells in a conditioned place preference (CPP) paradigm. **A**, Top, schematic of in vivo calcium imaging in hippocampal CA1 using a miniscope via an implanted GRIN lens. Bottom, illustration of the CPP box with the connecting door open. **B**, Left, schematic of the CPP experimental design. Right, longitudinally tracked place cell examples from Ctrl and MA mice. Each column is a cell. For each cell, the spatial firing rate map (warmer colors indicate higher firing rates) and raster plot (red dots) on top of animals running trajectory (black traces) are shown, organized according to the schematic on the left. **C,** Behavioral CPP scores (see Methods) for Ctrl and MA mice. MA mice showed a significant preference for the drug-paired side after the conditioning compared with the Ctrl in both test 1 (Ctrl vs. MA mean ± SEM: -5.62 ± 43.28 vs. 147.80 ± 37.21 s, t(15) = 2.68, p = 0.017, two-tailed unpaired t-test, n = 7 and 10, respectively) and test 2 (Ctrl vs. MA: 6.89 ± 28.67 vs. 204.10 ± 29.74 s, t(15) = 4.59, p = 0.0004, two-tailed unpaired t-test). The red-color labeled data point indicates a neutral effect of MA (defined as a negative CPP score) and data from this mouse was excluded from further analysis. **D**, Left, histology of an implanted GRIN lens above CA1. GCaMP6 expression shown in green, DAPI shown in blue. Right, enlarged view of GCaMP6-expressing CA1 pyramidal cells. **E,** Maximum intensity projected images of CA1 calcium imaging for baseline, test 1, and test 2 sessions from a representative mouse. Yellow arrows point to representative tracked neurons. Bottom right, CNMF-E spatial footprints of neurons extracted from the aligned and concatenated imaging data across all the sessions. **F,** Cytofluorogram of co-localization analysis between the baseline and test 2 images in (E). Fluorescence intensity for corresponding pixels of the two images in (E) are plotted, with warmer colors indicating higher data point frequency. The measured colocalization coefficient (Pearson’s) of 0.966 is higher than the 95^th^ % of the shuffled correlation coefficient (0.0062) shown on the right. See also Figure S1. **G,** Spatial correlation of each CPP compartment across sessions in Ctrl mice. Black bars show median and interquartile range of values for all place cells from all mice (median values: preferred side, bsl vs. test1 = 0.57, bsl vs. test2 = 0.58; non-preferred side, bsl vs. test1 = 0.56, bsl vs. test2 = 0.51; n = 1680 place cells from 7 mice), while colored circles indicate the mean value for each mouse. **H,** Left, Inter-compartment (between the left and right CPP compartment) rate map correlations of place cells in the baseline session for Ctrl (median: 0.46, n = 1680 place cells from 7 mice) and MA mice (median: 0.43, n = 2882 place cells from 9 MA mice). The red line (correlation = 0.4) indicates the threshold for separating global vs. rate remapped place cells. **I,** percentage change in peak Ca^2+^ event rates (peakER) between place fields from the two CPP compartments in baseline for rate remapped place cells. For each place cell, the value is defined as abs(peakER_left_ - peakER_right_) / max(peakER_left_, peakER_right_). Box plots show median, interquartile and full range of values for all the rate remapped place cells from all mice (median: Ctrl = 42.5%, MA = 43.2%, Z = -0.06, p = 0.95, two-tailed rank-sum test, n = 932 and 1556 rate remapped place cells, respectively), while colored circles indicate the mean value for each mouse.

We used Ai94; Camk2a-tTA; Camk2a-Cre transgenic mice (Figure 1D) to enable stable GCaMP6s expression in CA1 pyramidal neurons and single cell tracking across multiple days, as demonstrated in the maximum projected images and the colocalization analysis from different imaging sessions after alignments (Figures 1E and 1F, Figures S1A and S1B). We extracted calcium signals, using a CNMF based method (*53*), from aligned and concatenated imaged data for each animal, which facilitated maximum signal detection (*48, 54*) (Figure 1E). Calcium signals were then binarized into deconvolved spikes (*55*), which we treated as calcium events (Figure S1C).

In Ctrl animals, we observed stable spatial representations of CA1 place cells in both CPP compartments across days, with place cells defined by their activity in the baseline session (Figures 1B and 1G). Across the two CPP compartments, we observed signatures of both rate and global remapping in baseline sessions (Figures 1B and 1H). Ctrl and MA mice place cells showed similar inter-compartment (between the left and right CPP compartments) spatial correlation values in the baseline session, with ∼45% of place cells showing signatures of global remapping in both groups (defined as a correlation value less than 0.4) (Figure 1H). Between Ctrl and MA mice, place cells (∼55%) also showed similar levels of rate remapping in the baseline session, with a ∼43% change in peak calcium event rates between place fields in the two compartments (Figure 1I). Previous work has demonstrated that when animals are exposed to different novel environments, place cell representations become increasingly orthogonal between the environments as a function of experience (*37*). However, in Ctrl mice, spatial maps for the left versus right compartment in baseline did not show a decrease in similarity compared with the test sessions (median correlation: 0.43, 0.49, and 0.46, for baseline, test 1, and test 2 in Ctrl mice, respectively), suggesting the observed rate remapping in the baseline session is not due to insufficient experience in the CPP environment. Together these data demonstrate that in baseline sessions, CA1 place cells in both Ctrl and MA mice remap between the two contexts of a CPP paradigm.

### MA-conditioning results in a compartment specific decrease in place cell number

We next considered whether MA-conditioning induced changes in CA1 place cells at the population level. We calculated the Jaccard similarity between the population of place cells defined for each compartment and each session (pre-bsl, bsl, test1, and test2) (Figure 2A, Methods). The Jaccard similarity between two place cell populations measures the intersection of the two populations over the union, such that place cell representations with a high degree of overlap across sessions will show larger similarity values. In Ctrl mice, we did not observe a change in the Jaccard similarity of place cell representations between the baseline (bsl) and test sessions (test1 and test2) compared with the value obtained from the two baseline sessions (pre-bsl vs. bsl) (Figure 2A). However, in MA mice, we observed significant decreases in the Jaccard similarity between the baseline and test sessions preferentially on the saline-paired side of the apparatus (Figure 2A). To further investigate this decrease, we calculated the proportion of place cells over time for each animal in a compartment-specific manner. In MA mice, we found a significant decrease in the number of place cells on the saline-paired side in the test sessions compared with baseline (Figures 2B and 2C). In contrast, the number of place cells remained the same across sessions on both sides in Ctrl mice and on the MA-paired side in MA mice (Figures 2B and 2C). This indicates a selective decrease in the number of place cells in MA mice from baseline to test sessions on the saline-paired side of the CPP apparatus.

**Figure 2.**
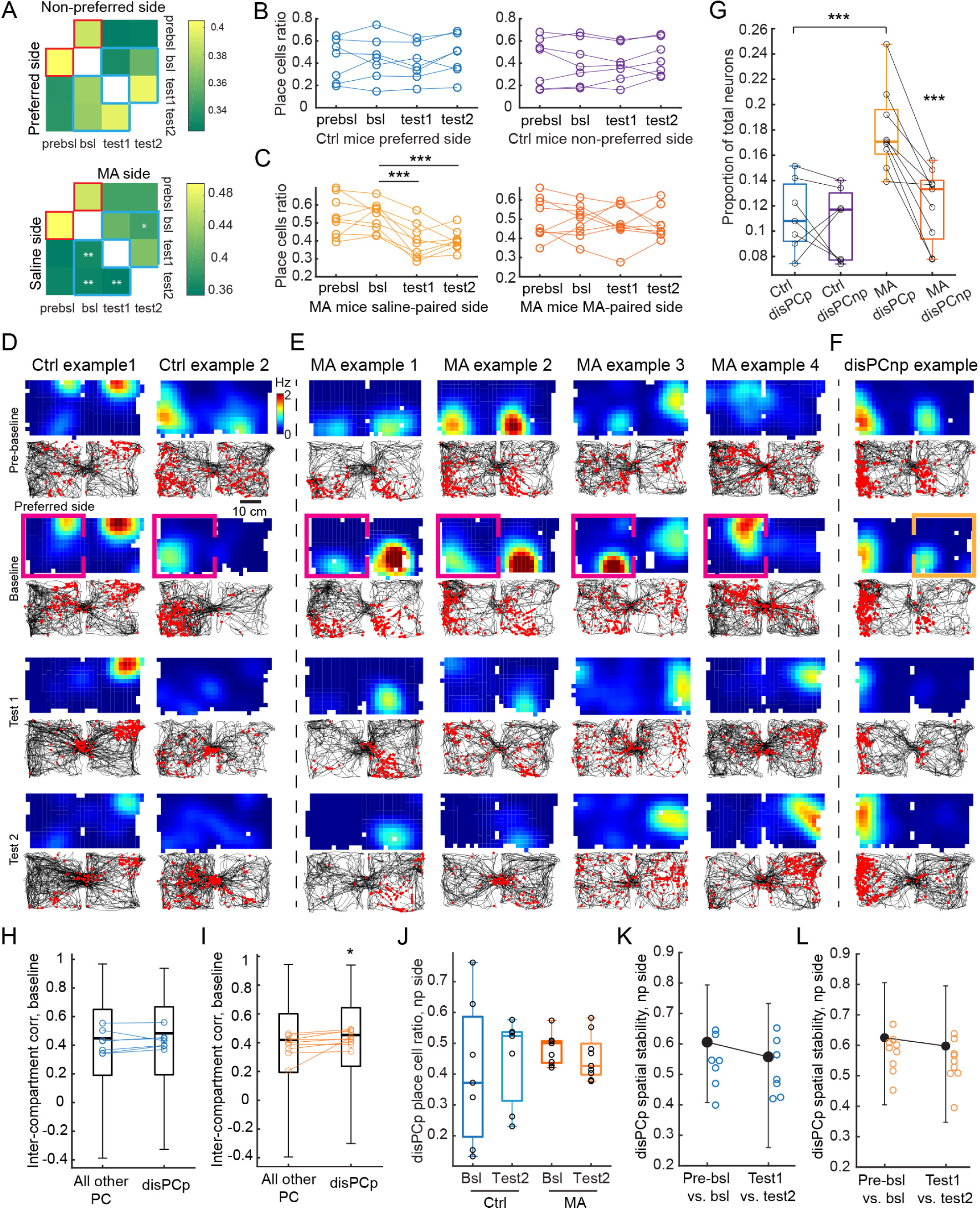
MA-conditioning results in a compartment specific decrease in place cell number. **A**, Jaccard similarity between populations of place cells in Ctrl (top) and MA (bottom) mice for any given two sessions. The lower triangle shows Jaccard similarity on the preferred/saline side, while the upper triangle shows the non-preferred/MA side. Statistical comparisons were made using the indices from bsl-test1, bsl-test2, and test1-test2 (outlined in blue) against the indices of bsl-prebsl (outlined in red) from the same side, respectively (** p < 0.01, * p < 0.05, two-tailed paired t-test, n = 9 mice). **B**, The ratio of place cells relative to the total number of neurons in Ctrl mice on the preferred and non-preferred side, respectively. **C**. Same as (B) but for MA mice. The number of saline-paired side place cells decreased in the test sessions (mean ± SEM, test 1: 0.39 ± 0.03, test 2: 0.40 ± 0.02,) compared to the baseline (0.53 ± 0.03) (bsl vs. test 1: t(8) = 6.30, p = 2.34 x 10^-4^; bsl vs. test 2: t(8) = 5.90, p = 3.64 x 10^-4^, two-tailed paired t-test, n = 9 mice). **D**, Two example disPCp from two different Ctrl mice. Each column is a cell tracked from baseline to test sessions. Top plots are rate maps with warmer colors indicating higher firing rates. Bottom plots show the mouse’s trajectory (black) and detected calcium events (red). Pink box indicates the session and compartment in which the place cell was defined. Plots are aligned such that the preferred side is on the left. **E**, Same as (D) but four example disPCp from four different MA mice. **F**, An example of disPCnp, orange box indicates the session and compartment in which the place cell was defined (non-preferred side). **G**, The proportion of disPCp and disPCnp relative to the total number of neurons in Ctrl and MA mice. In Ctrl mice, the proportion of disPCp (0.11 ± 0.01, mean ± SEM) was similar to disPCnp (0.11 ± 0.01) (t(6) = 0.57, p = 0.59, two-tailed paired t-test, n = 7). In MA mice, the proportion of disPCp (0.18 ± 0.01) was significantly higher than disPCnp (0.12 ± 0.01) (t(8) = 6.16, p = 2.7 x 10^-4^, two-tailed paired t-test, n = 9). The proportion of disPCp was significantly higher in MA compared to Ctrl mice (t(14) = 4.31, p = 0.00072, two-tailed unpaired t-test). The box plots show median, interquartile and full range of the data and black circles denote each individual data point. **H**, Inter-compartment rate map correlations for disPCp vs. all other place cells from baseline session in Ctrl mice (median: 0.43 vs. 0.43, respectively, p = 0.30, two-tailed sign-rank test, n = 7). The box plots show median, interquartile and full range of data from all cells while the colored circles show the mean values for each mouse. Statistic tests were performed based on circles. **I**, Same as (H) but for MA mice (median: 0.43 vs. 0.40, respectively; p = 0.012, two-tailed sign-rank test, n = 9). **J**, Place cell ratio of disPCp on the non-preferred/MA side. There was no change between baseline and test2 for either mouse group (Ctrl bsl vs. test: 0.41 ± 0.09 vs. 0.45 ± 0.05, mean ± SEM, t(6) = -0.61, p = 0.56, n = 7; MA bsl vs. test: 0.48 ± 0.02 vs. 0.45 ± 0.02, t(8) = 0.95, p = 0.37, N = 9; two-tailed paired t-test), nor the same session across the two groups (t(14) = -0.98, p = 0.34 for Ctrl bsl vs. MA bsl, t(14) = -0.12, p = 0.91 for Ctrl test vs. MA test, two-tailed unpaired t-test). The box plots show median, interquartile and full range of the data and black circles denote each individual data point. **K**, Spatial stability (Pearson’s correlation) of disPCp on the non-preferred side in Ctrl mice (F(1,568) = 1.40, p = 0.24, linear mixed effect model, n = 285 cells from 7 mice). **L**, Spatial stability of disPCp on the MA-paired side in MA mice (F(1,1358) = 1.72, p = 0.19, linear mixed effect model, n = 680 cells from 9 mice). For K and L, Black bars show median and interquartile range of values for all place cells from all mice, while colored circles indicate the mean value for each mouse.

To further analyze the decrease in place cells over time in MA mice, we identified place cells on the preferred (i.e., saline) side of the apparatus in the baseline session that lost their preferred/saline side spatial tuning in both test sessions (termed disPCp: <u>dis</u>appeared <u>p</u>lace <u>c</u>ells on the <u>p</u>referred side; note that in MA mice the preferred side is equivalent to the saline-paired side) (Figures 2D and 2E). Similarly, we identified place cells that lost their non-preferred (i.e., MA) side spatial tuning (termed disPCnp: <u>dis</u>appeared <u>p</u>lace <u>c</u>ells on the <u>n</u>on-<u>p</u>referred side) (Figure 2F). In MA mice, there was a greater proportion of disPCp than disPCnp, which indicates the place cell disruption we observed after MA conditioning was biased towards the preferred/saline side (Figure 2G). This effect was not observed in the Ctrl mice (Figure 2G). As expected, there was also a greater proportion of disPCp in MA compared to Ctrl mice (Figure 2G). Interestingly, while by definition the selection of disPCp is independent of their activity on the non-preferred/MA side, we found higher inter-compartment correlations for the baseline spatial maps of disPCp compared to all the other place cells in MA mice (Figure 2I). This was not observed in the Ctrl mice (Figure 2H). This suggests that MA has a larger effect on place cells with a high inter-compartment spatial correlation. To further investigate the spatial tuning of disPCp, we quantified the ratio of disPCp with a place field on the non-preferred side. We found no difference in this ratio before versus after MA conditioning in either MA or Ctrl mice (Figure 2J). In addition, MA conditioning did not affect the spatial stability of disPCp activity on the non-preferred side (Figures 2K and 2L). Together, these results demonstrate a selective disruption of preferred/saline side place fields in a subset of place cells after MA conditioning.

Finally, we employed a naïve Bayes classifier to examine whether the MA conditioning-induced changes to place cell numbers we observed impacted position coding. We trained the classifier on baseline data and then examined the accuracy of position decoding on data from test sessions (Figure S2A, Methods). We found that despite the larger number of disPCp in MA mice, Ctrl and MA mice showed comparable non-random overall decoding performance, as well as similar decoding accuracy between the preferred and non-preferred sides (Figures S2B - S2H). A principal component analysis revealed consistent results (Figures S2I and S2J). These data demonstrate that MA conditioning preferentially impacted the number of place cells on the preferred/saline CPP side without significantly altering the accuracy of an animal’s position coding.

### The activity of disPCp corresponds with encoding the MA-context association

As the larger number of disPCp in MA mice did not affect the accuracy of position coding (Figure S2), we next considered whether disPCp might contribute to the encoding of the MA-context association. To test this idea, we summed the spatial rate maps of all the disPCp to visualize their activity patterns (Figure 3A) and quantified an inter-compartment population vector (PV) correlation between corresponding spatial bins using all disPCp in a given mouse (Figure 3B, Methods). We found that the difference in the PV correlation (PV-diff) between the baseline and test sessions was significantly higher than zero in MA but not Ctrl mice (Figure 3A). Intuitively, a high PV-diff value indicates that the spatial maps of disPCp were less correlated between the two CPP compartments in the test compared to baseline session. Supporting the idea that disPCp may support the encoding of the MA-context, PV-diff was linearly correlated with an animal’s behavioral CPP score in MA but not Ctrl mice (Figure 3C). This indicates that, for MA mice, a higher CPP score corresponded to more orthogonalized inter-compartment spatial patterns in disPCp in the test versus baseline session.

**Figure 3.**
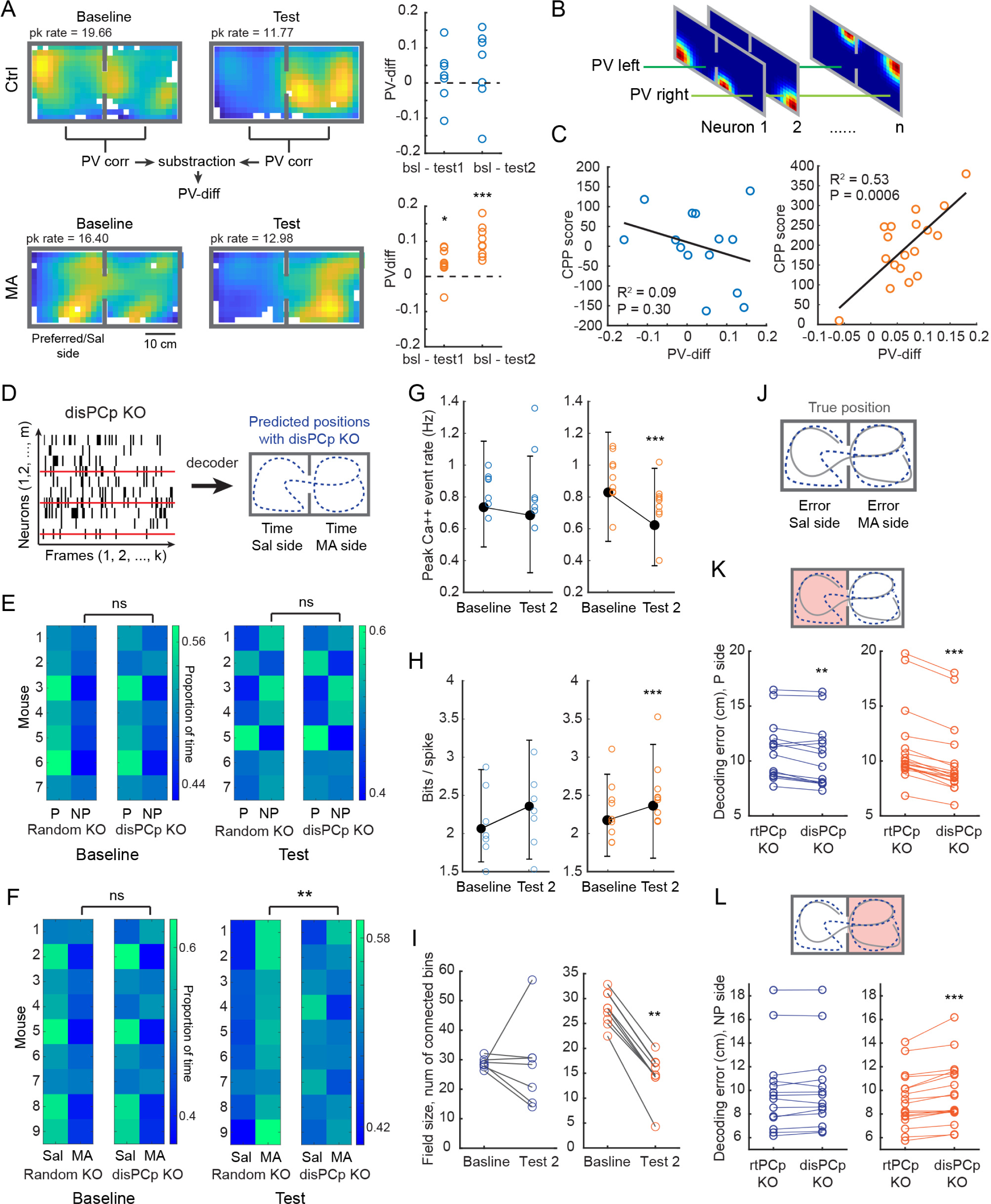
The activity of disPCp corresponds with the encoding of the MA-context association. **A**, Left, representative summed spatial rate maps of disPCp in baseline and test sessions for an example Ctrl and MA mouse. Warmer colors indicate higher firing rates. Right, PV-diff values for both Ctrl and MA mice. PV-diff values were not significantly different from zero in Ctrl mice (mean ± SEM, bsl – test1: 0.02 ± 0.03, t(6) = 0.71, p = 0.50; bsl – test2: 0.04 ± 0.04, t(6) = 1.06, p = 0.33; two-tailed t-test against zero, n = 7), but were significantly higher than zero in MA mice (bsl – test1: 0.038 ± 0.014, t(8) = 2.61, p = 0.031; bsl – test2: 0.099 ± 0.014, t(8) = 6.82, p = 1.35 x 10^-4^; two-tailed t-test against zero, n = 9). **B**, Schematic of population vectors (PV) in the CPP context. Inter-compartment PV correlations are the Pearson’s correlation between PV left and PV right, at the corresponding spatial bins. **C**, PV-diff was significantly correlated with behavioral CPP scores in MA (right) but not Ctrl mice (left). Each dot on the x-axis is a PV-diff value for each animal, as in (A). **D**, Schematic of the decoding method for the disPCp knock out (KO) analysis (see Methods). **E**, Reconstructed CPP time with disPCp vs. random neuron KO in Ctrl mice. Each row is a mouse and each column represents a CPP compartment (preferred/saline side is aligned on the left). Brighter colors indicate a larger proportion of total time. Left, baseline analyses (see results). disPCp KO (mean ± SEM, proportion of total time on the non-preferred side: 0.47 ± 0.01) did not lead to significant changes in the predicted CPP time compared with random KO (0.47 ± 0.01) (p = 0.08, two-tailed sign-rank test, n = 7). Right, test analyses. disPCp KO (0.50 ± 0.02) did not lead to significant changes in the predicted CPP time compared with random KO (0.51 ± 0.02) (p = 0.11, two-tailed sign-rank test, n = 7). **F**, Organized as in (E), for MA mice. Left, baseline analyses, disPCp KO (proportion of total time on the MA side: 0.46 ± 0.02) did not lead to significant changes in the predicted CPP time compared with random KO (0.44 ± 0.02) (p = 0.2, two-tailed sign-rank test, n = 9). Right, test analyses, disPCp KO (0.50 ± 0.01) significantly decreased the predicted CPP time on the MA-paired side compared with the random KO (0.55 ± 0.01) (p = 0.0039, two-tailed sign-rank test, n = 9). **G – H**, Place field properties of disPCp on the non-preferred/MA side (blue = Ctrl, orange = MA mice). In MA mice, MA conditioning decreased the peak firing rate (G, F(1,1358) = 14.06, p = 0.00018, linear mixed effect model, n = 680 cells from 9 mice) and increased spatial information (H, F(1,1358) = 18.17, p = 2.16 x 10^-5^) of disPCp. These changes were not observed in the Ctrl mice (peak firing rate: F(1,568) = 0.17, p = 0.68; spatial information: F(1,568) = 2.91, p = 0.09; linear mixed effect model, n = 285 cells from 7 mice). Black bars show median and interquartile range of values for all disPCp from all mice. Colored circles indicate the mean values for each animal. **I,** The field size of disPCp decreased on the non-preferred/MA side after conditioning in MA mice (median size bsl vs. test: 28.04 vs. 14.66 bins, p = 0.0039, two-tailed sign-rank test, n = 9), but not Ctrl mice (28.44 vs. 28.37 bins, p = 0.38, two-tailed sign-rank test, N = 7). **J**, Schematic for calculating compartment specific decoding errors. Grey line indicates true position and blue dotted line indicates decoded position. **K**, On the preferred/saline side, the decoding error for test sessions was larger after rtPCp KO compared to disPCp KO in both Ctrl and MA mice (mean ± SEM, rtPCp vs. disPCp in Ctrl: 10.99 ± 0.73 vs. 10.49 ± 0.78 cm, p = 0.0031; rtPCp vs. disPCp in MA: 11.11 ± 0.81 vs. 9.85 ± 0.76 cm, Z = 3.72, p = 1.96 x 10^-4^, two-tailed sign-rank test, n = 14 and 18 test sessions from Ctrl and MA mice). **L**, On the non-preferred/MA side, decoding error for test sessions was larger after disPCp KO compared to rtPCp KO in MA mice but not in Ctrl mice (rtPCp vs. disPCp in Ctrl: 9.95 ± 0.95 vs. 10.09 ± 0.94 cm, p = 0.09; rtPCp vs. disPCp in MA: 9.12 ± 0.54 vs. 9.72 ± 0.62 cm, Z = -3.59, p = 3.27 x 10^-4^, two-tailed sign-rank test).

To more directly consider the idea that disPCp contribute to the encoding of MA-context associations, we developed a knockout (KO) decoding analysis based on the aforementioned naïve Bayes classifier (Figure S2, Methods). In the KO decoding analyses, we ablated either disPCp or a random group of neurons (number matched) from the data set before feeding them into the trained classifier for making predictions (Figure 3D). We performed two sets of KO decoding analyses. First, we trained the decoder on one of the baseline sessions and then used the decoder to make the predictions on the other baseline session. Second, we trained the decoder on one of the test sessions and then used the decoder to make predictions on the other test session. To directly visualize the contribution of disPCp to CPP behavior, we plotted the reconstructed CPP time generated by the decoder for each individual mouse as heatmaps for each CPP compartment (Figures 3D - 3F). In Ctrl mice, there was no significant difference in the reconstructed CPP time for random versus disPCp KO in either the baseline or the test analysis (Figure 3E). Note, given the high decoding accuracy for both compartments (Figure S2), the results from the random KO condition largely recapitulated the true behavioral CPP time, with more reconstructed time shown on the preferred side (left column of Figures 3E and 3F). In MA mice, there was no significant difference in the reconstructed CPP time for random versus disPCp KO in the baseline analysis (Figure 3F, left). Strikingly however, in MA mice, the reconstructed CPP time clearly captured the MA-induced place preference for the random KO but this place preference was disrupted for the disPCp KO (Figure 3F, right). Together these results reveal that in MA mice, disPCp shift from encoding the two CPP compartments relatively equally (Figure 3F) to differentially encoding the two compartments after the MA conditioning.

### The activity of disPCp sharpens on the MA-paired side after MA conditioning

As disPCp place fields were disrupted on the preferred/saline side in the test sessions, their activity on the non-preferred/MA side could indicate the function of disPCp after MA conditioning. Thus, we next examined disPCp place cell metrics for place fields on the non-preferred/MA side. In MA mice, we found disPCp place fields on the non-preferred/MA side had a decreased peak calcium event rate, increased spatial information and a smaller field size between the baseline and test2 session (Figures 3G - 3I). In contrast, in Ctrl mice, disPCp place fields on the non-preferred side did not differ in their peak calcium event rate, spatial information or field size between the baseline and test2 session (Figures 3G - 3I). These results suggest that disPCp strengthened their coding for the MA-paired side, in addition to losing their spatial tuning on the preferred/saline side after MA conditioning.

Next, we considered how much the sharpening of disPCp activity contributed to the encoding of MA-context associations, compared to other place cells. For this analysis, we considered place cells with place fields on the preferred/saline side in baseline that also retained their spatial tuning in at least one of the test sessions (termed rtPCp: retained <u>p</u>lace <u>c</u>ells on the <u>p</u>referred side. Note disPCp and rtPCp are mutually exclusive). We then compared the contribution of disPCp and rtPCp to spatial coding of the non-preferred/MA side after conditioning. We again trained the naïve Bayes decoder on data from the baseline session and then examined the accuracy of position decoding on data from the test sessions. We performed two sets of decoding analyses, ‘knocking-out’ either disPCp or rtPCp (number matched) from the test sessions data, and then compared decoding error within each CPP compartment (Figures 3D and 3J, Methods). Intuitively, a smaller decoding error indicates the ablated neurons become less spatially informative after conditioning. We found that disPCp KO resulted in smaller decoding errors compared to rtPCp KO on the preferred/saline side in both the Ctrl and MA mice (Figure 3K), as the disPCp were no longer place cells on the preferred/saline side in the test sessions and thus carried less information than the rtPCp regarding the preferred side context. In contrast, on the non-preferred/MA side, rtPCp and disPCp KO showed comparable decoding error in Ctrl mice (Figure 3L). However, in MA mice, disPCp KO resulted in significantly larger decoding errors than rtPCp KO (Figure 3L). This result suggests disPCp contribute more to the spatial coding of the MA-paired context than rtPCp after MA conditioning. This points to the disPCp as the primary contributor to the encoding of MA-context associations.

### disPCp in MA mice emerge in an experience-dependent manner

To further explore the possible link between disPCp and the encoding of context during drug-conditioning, we asked whether the spatial location of disPCp place fields was biased towards locations mice had physically visited during conditioning sessions. In the CPP paradigm, traversals between the two compartments are possible during the baseline and test sessions but are not possible during the conditioning sessions (Figure 4A). In baseline sessions, we observed a sub-set of place cells fired specifically at the junction between the two CPP compartments (‘center-firing’ neurons) (Figure 4B). As inter-compartment traversals were not possible during the conditioning sessions, we hypothesized that if the disPCp specifically encode the MA-associated contextual experience, the disPCp population should contain very few ‘center-firing’ neurons.

**Figure 4.**
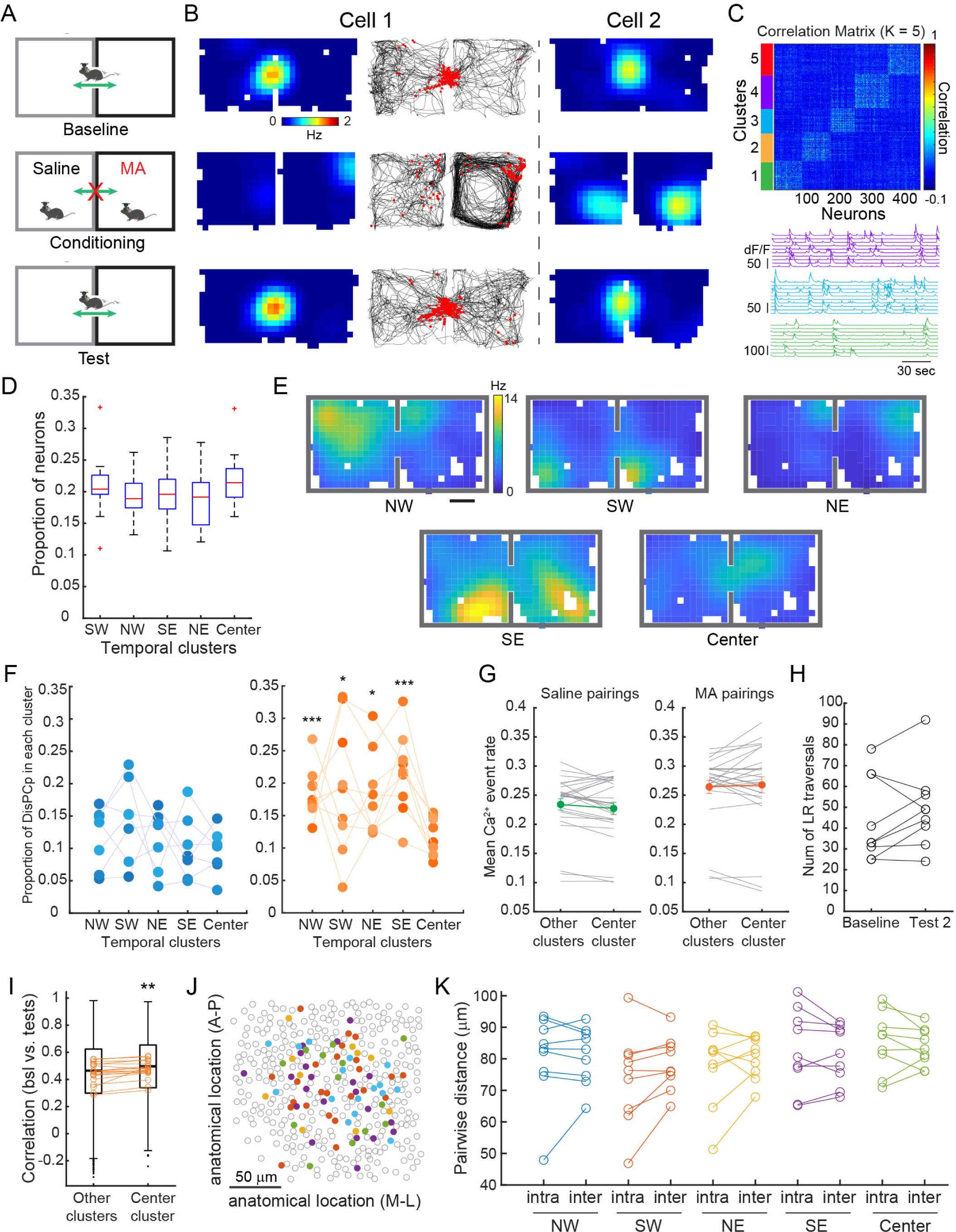
disPCp in MA mice emerged in an experience-dependent manner. A, Schematic illustrating an animal’s inter-compartment movement in baseline, conditioning, and test sessions. **B,** Two example neurons that are maximally active at the junction between the two compartments. Session organized as labeled in (A). Left, rate map (warmer colors indicate higher firing rates) and raster plot (red calcium events on top of black running trajectory) from one neuron. Right, rate map of a neuron from a different mouse **C,** Top, the temporal correlation matrix of calcium signals from the baseline session of an example mouse. The matrix was sorted by the k-means derived temporal clusters. Bottom, 10 representative calcium traces from clusters 4, 3, and 1 (colors correspond to the top panel) illustrating the synchronized calcium activity within each cluster. Each line corresponds to an individual neuron. **D,** For Ctrl and MA mice, the proportion of neurons assigned to each cluster was roughly equal (F(4,75) = 1.77, p = 0.14, one-way ANOVA, n = 16 mice from Ctrl and MA groups). The box plots show median, interquartile and full range of the data without outliers. The red ’+’ symbol indicates outliers. SW, southwest; NW, northwest; SE, southeast; NE, northeast. **E**, Summed ensemble rate maps of temporally clustered disPCp from an MA mouse. **F,** disPCp showed a non-random distribution across temporal clusters in MA (orange) but not Ctrl (blue) mice (F(4,30) = 1.06, p = 0.39 for Ctrl; F(4,40) = 3.01, p = 0.029 for MA, one-way ANOVA, n = 7 and 9, respectively). In MA mice, the proportion of disPCp in the center cluster (mean ± SEM, 0.12 ± 0.01) was significantly lower compared to other clusters (NW: 0.19 ± 0.01, SW: 0.19 ± 0.03, NE: 0.18 ± 0.02, SE: 0.22 ± 0.02; NW vs. center, t(8) = 5.87, p = 1.84 x 10^-4^; SW vs. center, t(8) = 1.95, p = 0.043; NE vs. center, t(8) = 2.35, p = 0.023; SE vs. center, t(8) = 5.33, p = 3.50 x 10^-4^; one-tailed paired t-test, n = 9). **G**, The activity of neurons from center vs. other clusters in saline or MA conditioning sessions for MA mice (Z = 1.54, p = 0.12 and Z = -0.41, p = 0.68, two-tailed sign-rank test, n = 27 conditioning sessions from 9 mice). Gray lines represent individual conditioning sessions, colored lines indicate the mean ± SEM (left, 0.23 ± 0.01 vs. 0.23 ± 0.01; right, 0.26 ± 0.01 vs. 0.27 ± 0.01). **H,** The number of inter-compartment traversals between the baseline and test 2 in MA mice (mean ± SEM, 44.22 ± 6.73 vs. 49.44 ± 6.43, t(8) = -1.29, p = 0.23, two-tailed paired t-test, n = 9 mice). **I**, Pearson’s correlation of rate maps between baseline and test sessions from neurons in center vs. other clusters in MA mice. The box plots show correlation values (baseline vs. test 1 and baseline vs. test 2) from all neurons in center (n = 740 cells) vs. other clusters (n = 3143 cells), black dots are outliers. The colored circles show the mean values for each mouse (n = 18 session comparisons from 9 mice). Center clustered neurons show significant higher correlation than other clustered neurons (p = 0.0012, Z = 3.24, two-tailed sign-rank test, statistic test is based on the circle-labeled mean values). **J**, Anatomical location of disPCp from a representative mouse. Each filled dot is a centroid of one disPCp neuron color coded for different temporal clusters. Grey circles are the centroids of other recorded CA1 neurons that were not disPCp. **K,** There was no difference between the pairwise intra- vs. inter-cluster anatomical distances for any disPCp cluster in MA mice (intra vs. inter: mean ± SEM (μm), NW: 80.43 ± 4.62 vs. 81.17 ± 3.04, t(8) = -0.34, p = 0.74; SW: 73.79 ± 4.95 vs. 78.43 ± 2.85, t(8) = -1.97, p = 0.08; NE: 77.55 ± 4.13 vs. 80.16 ± 2.12, t(8) = -0.95, p = 0.37; SE: 82.77 ± 4.29 vs. 81.39 ± 3.06, t(8) = 0.83, p = 0.43; Center: 85.42 ± 3.19 vs. 83.76 ± 1.98, t(8) = 0.88, p = 0.40; two-tailed paired t-test, n = 9).

To classify neurons as ‘center-firing’, we used a consensus k-means method to group neurons into temporally synchronized clusters using the temporal correlations of their calcium signals in the baseline session (Figure 4C, Methods). We determined the optimal number of clusters (k = 5) using two different measurements (Figure S3, Methods). This clustering approach both identified the ‘center cluster’ at the junction and grouped the remaining neurons into four clusters with peak ensemble activity at different spatial positions within the CPP compartments (Northeast [NE], Northwest [NW], Southeast [SE] and Southwest [SW]) (Figure S4). The temporal clustering pattern was consistent across animals, which enabled us to use a template matching method to sort k-means derived temporal clusters into five clusters according to the location of their ensemble activity peaks (Figure S4). As place cells are spatially tuned, we also performed spatial clustering using the baseline spatial tuning of neurons and found compared to spatial clustering, temporal clustering was better at capturing the firing pattern of neurons at the junction (Figure S5).

The proportion of total neurons assigned to each cluster were roughly equal across animals (Figure 4D). However, while the proportion of disPCp was equally distributed across clusters in Ctrl mice, it was not equally distributed across clusters in MA mice (Ctrl p = 0.39, MA p = 0.029, one-way ANOVA) (Figures 4E and 4F). For MA mice, we visualized this unequal distribution of the ensemble activity by plotting the summed disPCp rate maps from each cluster (Figure 4E, from an example mouse). Further quantification of the disPCp distribution across clusters in MA mice revealed that, while there was no difference in the proportion of disPCp across the NW, SW, NE and SE clusters, the center cluster contained a smaller proportion of disPCp compared to all other clusters (Figure 4F). The smaller proportion of disPCp in the center clustered neurons was not due to insufficient neural activity during the conditioning sessions in MA mice, as center clustered neurons often remapped to represent a different spatial location during the conditioning session (Figure 4B), and we observed a comparable level of mean Ca^2+^ event rates between center clustered neurons and neurons from other clusters (Figure 4G). In addition, we did not observe any differences in the number of traversals across the junction of the CPP compartments (Figure 4H) or the running speed between baseline and test sessions (Figure S6C). However, compared to all other clusters, center-clustered neurons showed a higher spatial map correlation between the baseline and test sessions of MA mice (Figure 4I). This indicates center clustered neurons exhibited more stable spatial representations from baseline to test, regardless of the influence of repeated MA conditioning (Figures 4B and 4I). Interestingly, in MA mice, the anatomical distribution across CA1 of each disPCp cluster was random (Figure 4J), as demonstrated by the quantification of disPCp pairwise intra- vs. inter-cluster distances (Figure 4K). Together, these results suggest that disPCp emerge in an experience-dependent manner and further strengthen the link between disPCp and the encoding of context during drug-conditioning.

### Ketamine disrupts MA CPP and the neural signatures of drug-context associative learning

We next aimed to investigate whether the involvement of disPCp is dependent on the intact drug-context experience. We considered whether we could disrupt MA CPP by dissociating the drug-induced affective state from sensory inputs and examined what impact this disruption might have on CA1 place cells. To achieve this dissociation state, we used a subanesthetic dosage (25 mg/kg intraperitoneal) of ketamine, which has been shown to induce dissociation-like states in mice by disconnecting the sensory motor function from affective and motivational responses (*52*). To test ketamine’s effect on MA CPP, and avoid confounding the potentially addictive nature of ketamine, we paired ketamine with both saline and MA during the conditioning sessions (Figure 5A).

**Figure 5.**
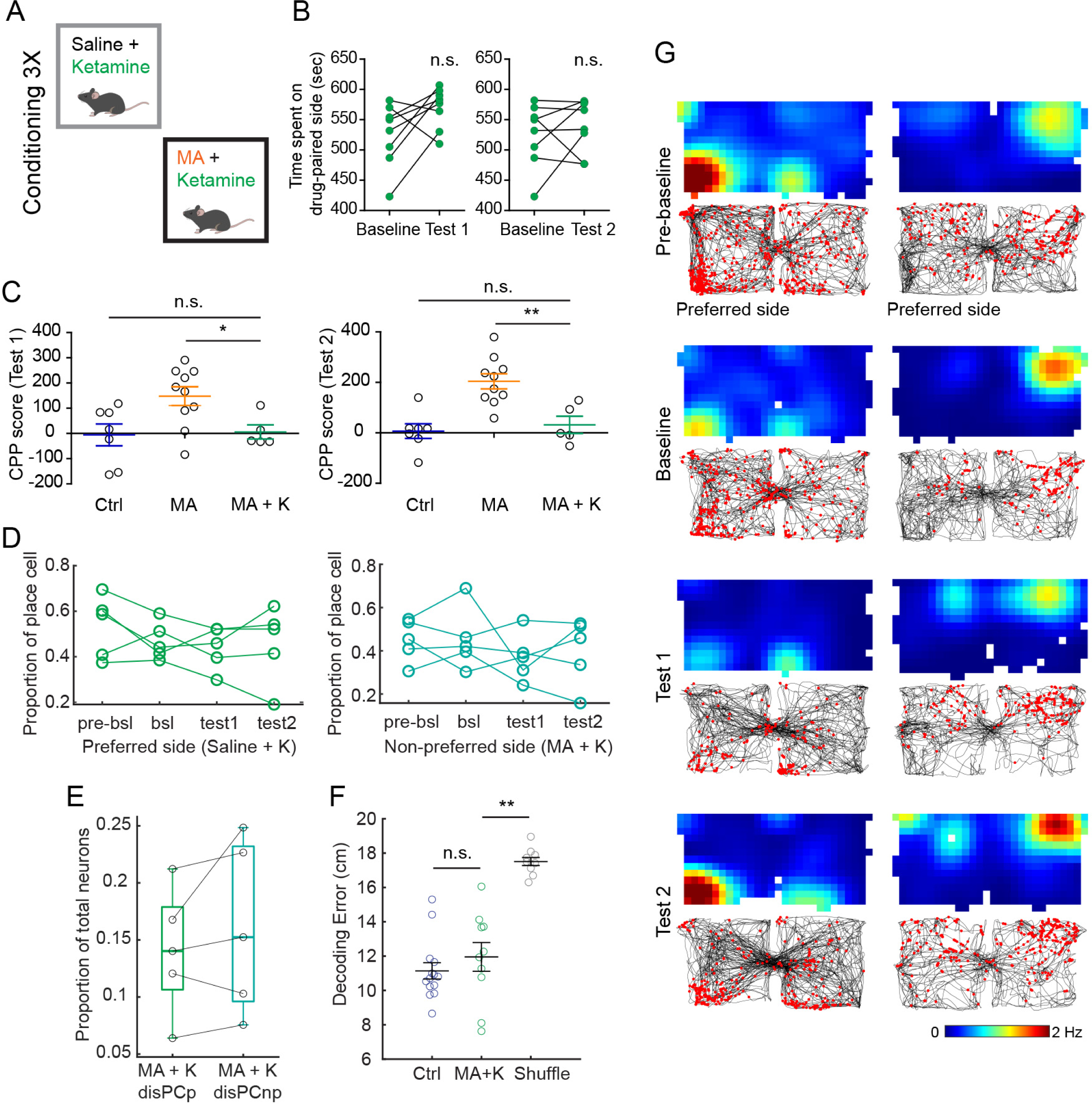
Ketamine disrupts MA CPP and the neural signatures of drug-context associative learning. **A**, Experimental design for the conditioning sessions of MA CPP with ketamine (MA+K). **B**, Behavior results of MA CPP with ketamine in wild type C57BL/6 mice. The time that animals spent on the MA-paired side in test sessions (mean ± SEM: 569.9 ± 11.96 and 539.9 ± 15.45 s for test 1 and test 2, respectively) was not different from the baseline (525.4 ± 18.44 s)(bsl vs. test1: t(7) = 2.08, p = 0.076; bsl vs. test 2: t(7) = 0.74, p = 0.48; two-tailed paired t-test, n = 8 mice). **C**, CPP score for imaged mice. Ketamine significantly decreased MA-induced place preference in MA+K mice compared with MA mice for both the test 1 (Ctrl: -5.62 ± 43.68 s, mean ± SEM, MA: 147.8 ± 37.21 s, MA+K: MA: 5.9 ± 27.66 s) and test 2 (Ctrl: 6.89 ± 28.67 s, MA: 204.1 ± 29.74 s, MA+K: MA: 31.85 ± 33.91 s). Statistic tests for test 1: Ctrl (n = 7) vs. MA+K (n = 5), t(10) = 0.20, p = 0.84; MA (n = 10) vs. MA+K, t(13) = 2.50, p = 0.027. Statistic tests for test 2: Ctrl vs. MA+K, t(10) = 0.56, p = 0.59; MA vs. MA+K, t(13) = 3.54, p = 0.0036. All statistic tests were two-tailed unpaired t-tests. **D**, In MA+K mice, there was no significant decrease in the proportion of place cells in test sessions compared with baseline on the preferred side (bsl: 0.47 ± 0.04, mean ± SEM, test1: 0.44 ± 0.04, test2: 0.46 ± 0.07; bsl vs. test1, t(4) = 0.73, p = 0.51; bsl vs. test 2, t(4) = 0.15, p = 0.89; two tailed paired t-test, n = 5 mice) or non-preferred side (bsl: 0.45 ± 0.06, test1: 0.37 ± 0.05, test2: 0.40 ± 0.07; bsl vs. test1, t(4) = 0.97, p = 0.39; bsl vs. test 2, t(4) = 0.75, p = 0.50; two-tailed paired t-test, n = 5). **E**, There was no significant difference between the proportion of disPCp and disPCnp over the total number of neurons in MA+K mice (mean ± SEM: 0.14 ± 0.02 vs. 0.16 ± 0.03, t(4) = -1.25, p = 0.28, two-tailed paired t-test, n = 5). The box plots show median, interquartile and full range of the data and black circles denote each individual data point. **F**, Decoding error of Ctrl, MA+K, and MA+K shuffle conditions in test sessions (test 1 and 2) using a baseline trained decoder. Decoding error in MA+K mice (mean ± SEM: 11.95 ± 0.84 cm) was similar to Ctrl (11.14 ± 0.47 cm) and significantly lower than the shuffled condition (17.51 ± 0.23 cm). Statistic tests: Ctrl vs. MA+K, Z = -1.2, p = 0.23, two-tailed rank sum test, n = 14 test sessions from 7 mice and 10 test sessions from 5 mice, respectively. MA+K vs. MA+K shuffle, p = 0.002, two-tailed sign-rank test, n = 10 test sessions from 5 mice. **G**, Rate maps (top, warmer colors indicate higher firing rates) and raster plots (bottom, red calcium events on top of black running trajectory) for two example place cells are shown from two different MA+K mice.

We first performed MA CPP paired with ketamine (Figure 5A) in a cohort of wild type C57BL/6 mice (n = 8). There were no significant differences between either test session vs. baseline in the time the animals spent on the MA-paired side (Figure 5B). We then performed miniscope imaging using the same CPP design in a cohort of Ai94;Camk2a-tTa;Camk2a-Cre mice (N = 5; group name: MA+K). Consistent with wild type mice, cross group comparisons in MA+K mice demonstrated that ketamine significantly decreased the CPP score in both test sessions (Figure 5C). In addition, the proportion of place cells on each CPP side in MA+K mice did not significantly decrease between the baseline and test sessions and the proportion of disPCp compared with disPCnp did not significantly change (Figures 5D and 5E). Furthermore, MA conditioning with ketamine did not affect the long-term spatial coding accuracy of neurons in test compared to baseline sessions, as indicated by the accuracy of position estimates in the test sessions using a decoder trained on the baseline session and the stability of place field locations (Figures 5F and 5G). Together, these data demonstrate that ketamine blocks MA CPP and provide further evidence that the presence of disPCp requires an intact MA-context experience.

### disPCp encode morphine (MO)-associated spatial context

We next considered whether the coding features of disPCp in drug-context associations generalize to other addictive substances. To address this question, we performed a morphine (MO) CPP experiment that followed the same protocol as the MA CPP. As in MA mice, we observed stereotyped behavior during MO conditioning (Figure S7A, Figure 1B) and significant place preference for the MO-paired compartment in the test sessions (Figure S7B). Similarly, as with MA conditioning, MO conditioning did not significantly alter the overall spatial coding accuracy in the test sessions using a baseline trained decoder (Figures S7C - S7E). Consistent with our observations in MA mice, place cell numbers decreased specifically on the preferred/saline-paired side in MO mice (Figure 6A) and, as in MA mice, this decrease reflected a greater proportion of disPCp compared with the disPCnp (Figure 6B). Interestingly, as in MA mice, PV-diff of disPCp in MO mice significantly correlated with the CPP score but, in contrast, this correlation was negative in MO mice (Figure 6C). In addition, while MA disPCp equally encoded the two CPP compartments in baseline (Figure 3F), MO disPCp preferentially encoded the preferred/saline side in baseline (Figure 6D, left). Consistent with this observation, the proportion of MO disPCp that expressed a place field on the non-preferred side in the baseline was significantly lower than that in MA disPCp (Figures 6E and 6G). Moreover, the comparison of PV correlation in baseline suggested MA and MO disPCp were two different place cells populations in terms of their inter-compartmental spatial firing patterns (Figure 6F), with MO disPCp showing more orthogonal firing patterns across the two compartments (Figures 6F and 6G). However, like disPCp in MA mice, disPCp in MO mice biasedly encoded the non-preferred/MO side after conditioning (Figure 6D, right). Even so, unlike MA disPCp, the place fields of MO disPCp on the MO-paired side did not show any change in spatial information or place field size after MO conditioning (Figures S7F and S7G). Together, these data suggest MO also engages disPCp to encode contextual associations but preferentially recruits place cells that show a biased spatial coding towards the preferred/saline side in baseline and shifts their spatial coding to the MO-paired side after conditioning.

**Figure 6.**
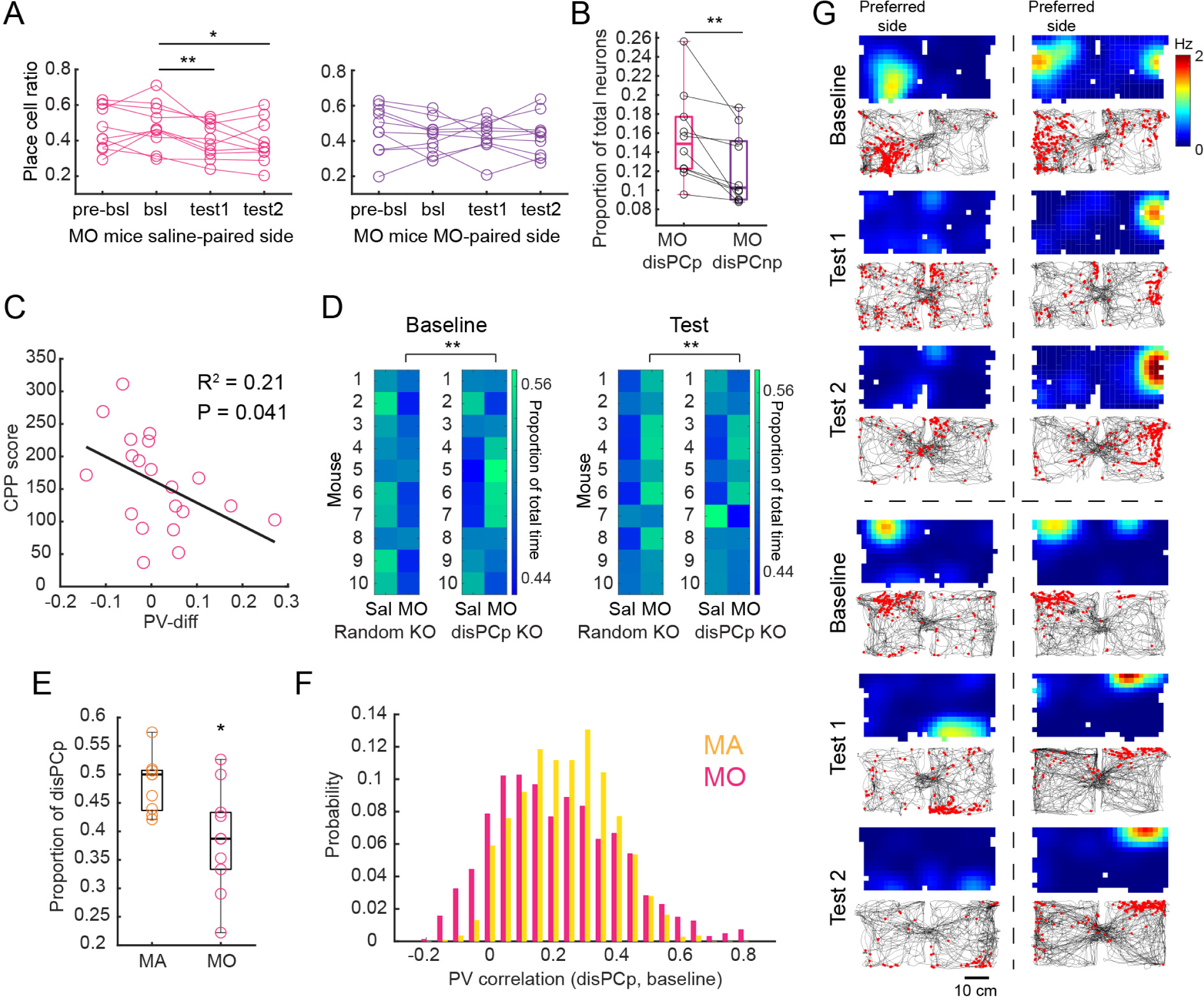
disPCp encode morphine (MO)-associated spatial context. **A**, The ratio of place cells relative to the total number of neurons in MO mice on the saline-paired and MO-paired side, respectively. The number of saline-paired side place cells decreased in the test sessions (mean ± SEM, test 1: 0.40 ± 0.03, test 2: 0.39 ± 0.04) compared to the baseline (0.48 ± 0.04) (bsl vs. test 1: t(9) = 3.5, p = 0.0067; bsl vs. test 2: t(9) = 2.50, p = 0.034, two-tailed paired t-test, n = 10). **B**, In MO mice, the proportion of disPCp (mean ± SEM: 0.16 ± 0.01) was significantly higher than disPCnp (0.12 ± 0.01) (t(9) = 3.32, p = 0.0090, two-tailed paired t-test, n = 10). The box plots show median, interquartile and full range of the data and the black circles denote each individual data point. **C**, PV-diff was significantly correlated with the CPP scores in MO mice. Each circle in the plot is a session difference for each animal (baseline – test 1 or baseline – test 2, n = 20 session differences from 10 mice). **D**, Reconstructed CPP time based on the decoding prediction with disPCp vs. random neuron KO in MO mice. Each row is a mouse and each column represents a CPP compartment (preferred/saline side is aligned on the left). Brighter colors indicate a larger proportion of total time. Left, for baseline analysis, disPCp KO (proportion of total time on the MO side: 0.51 ± 0.01, mean ± SEM) significantly increased the predicted CPP time on the non- preferred/MO side compared with random KO (0.48 ± 0.007) (p = 0.0039, two-tailed sign-rank test, n = 10). Right, for test analysis, disPCp KO (0.50 ± 0.01) significantly decreased the predicted CPP time on the MO-paired side compared with random KO (0.53 ± 0.007) (p = 0.0019, two-tailed sign-rank test, n = 7). **E**, In the baseline session, the proportion of MO disPCp that expresses a place field on the non-preferred side is significantly lower than that in MA disPCp (mean ± SEM, MO: 0.38 ± 0.03, n = 10, MA: 0.48 ± 0.02, n = 9; t(17) = 2.80, p = 0.012, two-tailed unpaired t- test). The box plots show median, interquartile and full range of the data and the colored circles denote each individual data point. **F**, Histograms showing inter-compartment PV correlations of disPCp in the baseline session from MA and MO mice, respectively. The histogram pulls all the PV correlations for each corresponding spatial bin from all the mice in each group. (MA: median = 0.24, n = 1792 PV correlations from 9 mice; MO: median = 0.19, n = 2016 PV correlations from 10 mice). The two distributions are significantly different from each other (p = 8.89 x 10^-14^, z = 0.14, two-sample Kolmogorov-Smirnov test). **G**, Rate maps (top, warmer colors indicate higher firing rates) and raster plots (bottom, red calcium events on top of black running trajectory) of four disPCp examples are shown from three different MO mice. The top two examples are from the same mouse.

Given both MA and MO mice use disPCp to encode drug-context association, the differential mechanisms of disPCp may result from the different drug-induced affective states. Thus, we hypothesized MA and MO should exert differential direct impacts on place cell representation. To test this hypothesis, we performed separate decoding analysis for conditioning sessions by training the naïve Bayes classifiers using either the preferred or non-preferred side data from baseline and making predictions using the conditioning data from the corresponding side (Figure S8A). By comparing decoding errors from the first conditioning session across groups, we found although mice from different groups showed similar decoding performance during the first saline pairing (Figures S8B and S8C), decoding errors of MO mice in the first drug pairing were significantly higher than both Ctrl and MA mice (Figures S8B and S8D). This result suggests there is a significant remapping of place field locations upon the first exposure to MO but not MA (Figure S8B). By comparing decoding errors between the first and last conditioning sessions within each group, we found, in MA mice, decoding errors from the last conditioning session were significantly greater than the first, in both the saline- and MA-pairing (Figures S8E and S8F). In contrast, MO mice showed comparable decoding errors across time on both sides (Figures S8E and S8F). Together, these results suggest MA induces a progressive place cell remapping by moving the place fields on both sides gradually away from their original location along with repeated drug experience (Figure S8B). However, MO induces a direct location remapping upon the first drug exposure and maintains this remapping status in the following drug experience without affecting the saline-paired side (Figure S8B).

## Discussion

The coding of spatial positions by CA1 place cells, as well as the sensitivity of place cell coding features to reward, points to the hippocampus as a potential candidate brain region for the pairing of a spatial environment with the rewarding experience of drug administration. Consistent with this idea, our experiments revealed a subset of CA1 place cells (disPCp) that altered their coding features over the course of drug-context associative learning. During MA conditioning, place cells that rate remapped between the two CPP compartments were more likely to selectively encode the MA-paired compartment post-conditioning. In comparison, during MO conditioning, place cells that globally remapped between the two CPP compartments were more likely to selectively encode the MO-paired compartment post-conditioning. In both MA and MO conditioning, we observed a significant correlation between the change of compartment selectivity of disPCp and a mouse’s behavioral post-conditioning preference for the drug-paired CPP context. Further raising the possibility of a link between disPCp and CPP behavior, ketamine co-administered with MA blocked both the post-conditioning emergence of disPCp and CPP behavior. Together, this work reveals that a sub-population of hippocampal CA1 place cells encode drug-associated contextual information, pointing to a potential novel neuronal target for the development of therapeutics for treating context-induced drug relapse (*56–58*).

Previous works have shown that place cell activity can be modulated by spatial locations with a high behavioral significance, such as goal locations associated with rewards (e.g., food or water). Several phenomena have been observed regarding reward-driven changes to hippocampal place cell coding features. First, place cell firing fields often accumulate near locations associated with reward (*45, 47, 59, 60*). The over-representation of reward locations by place cells typically develops through experience, with the firing field of place cells gradually shifting closer to the reward or goal location over behavioral trials or sessions (*35, 36, 61-63*). Moreover, this over-representation of reward locations also reflects the activity of a small population of hippocampal neurons dedicated to coding reward (*59*). Second, studies have reported elevated activity of existing place cells (i.e., out-of-field firing activity) around reward or goal locations, which is not accompanied by an accumulation of place fields (*64–66*). Our results differ from these previous works, in that we did not observe shifting of place fields from the saline- to the drug-paired context nor did we observe an increase in firing activity in the drug-paired context, which could reflect a differential effect of natural rewards and addictive drugs on hippocampal place cells. However, the loss of disPCp place fields selectively in the saline-paired context does increase the relative representation of the drug-paired context by the hippocampal place cell population. This finding is broadly consistent with previous observations of the over- representation of rewarded locations by place cells and in line with the notion that addictive drugs may usurp the normal neural machinery for learning about reward or goal locations to achieve a similar enduring association with specific spatial locations (*67–69*).

Our observation of similar firing patterns in place cells across the two CPP compartments in baseline is also consistent with previous works have shown that place cells can exhibit similar place field locations across geometrically identical contexts or segments of linear tracks (*70–73*). Under such conditions however, place cells often rate remap, which could support firing rate-based discrimination of contexts. Interestingly, in the current work, MA-conditioning disPCp were preferentially place cells that rate remapped between the two CPP compartments in baseline. One possibility is that place cells that rate remap between the compartments are primed to differentially encode features that generalize across the two contexts, with this differential encoding amplified by the pairing of MA with one of the contexts. Such amplification could then result in better pattern separation between the two CPP compartments, which could drive stronger memory encoding and recall of the pairing between the drug and spatial context. This idea is broadly consistent with our observations of place cells during MO-conditioning, in which MO- conditioning disPCp were preferentially place cells that globally remapped between the two CPP compartments in baseline. Place cells that globally remap between compartments may be primed to differentially encode features that are distinct between the two contexts. In this case, the preferential effect of MO on place cells that globally remap may allow MO to form a stronger association with contextual features that are highly specific to the drug context. One possible interpretation then, is that features in the environment may be differentially associated with different addictive drugs. An alternative interpretation is that the neural representation changes in disPCp reflect a value-based signal that facilitates a comparison between drug versus saline-paired contexts. However, previous works have found little evidence for the influence of value on hippocampal neural coding, (*65, 74–76*). Thus a more likely interpretation is that the disPCp provide a mechanism for encoding drug-associated contextual information, which then goes on to inform value coding in other brain regions and drive drug seeking behavior (*77, 78*).

Our observations of MA and MO driven changes to hippocampal place cell representations almost certainly interface with other brain regions to encode drug-associated contexts and drive drug-seeking behavior. Given the biochemical nature of MA and MO, direct monoamine inputs from locus coeruleus (*45, 79, 80*), VTA (*81–83*), and raphe nuclei (*84, 85*) could all play a role in the emergence of disPCp. Hippocampal representations of drug-associated contexts then likely interface with regions like the nucleus accumbens (NAc), one of the major efferent targets of the hippocampal formation, with hippocampal-NAc circuit interactions playing a key role in spatial context conditioning (*10, 22, 86*). Notably, recent work revealed that cocaine place conditioning increases the functional drive from hippocampal CA1 neurons that encode cocaine-paired locations to NAc medium spiny neurons (*18*). Thus, one possibility is that the disPCp route information directly to the NAc to encode drug-associated spatial contexts, with this circuit then potentially provoking future drug-seeking behavior (*87*). Alternatively, disPCp may route information to the subiculum (Groenewegen et al., 1987), where electrical stimulation has been shown to increase dopamine levels in NAc and trigger cocaine relapse that resembles context-induced drug reinstatement (*21, 83*). Another potential efferent target is the medial entorhinal cortex, which innervates the NAc (Ohara et al., 2018; Sürmeli et al., 2015) and contains neurons that both encode the position and orientation of an animal that are modulated by reward locations (*88, 89*). Further investigation will be needed to identify the full circuit mechanisms by which disPCp interact with known reward circuitry to drive drug-seeking behavior.

The question remains as to the mechanism underlying the preferential recruitment of rate remapping place cells as disPCp during MA-conditioning versus global remapping place cells as disPCp during MO-conditioning. This question intersects with the differential correlation of disPCp with MA- versus MO-conditioned CPP behavior: the orthogonalization of disPCp positively correlated with MA-conditioned CPP behavior while the orthogonalization of disPCp negatively correlated with MO-conditioned CPP behavior. One possibility is that, unlike psychostimulants, opioid usage produces withdraw effects that contribute to subsequent compulsive drug taking (*69, 90*). However, we did not observe a change in the time the animal’s spent in the saline-paired compartment over MO conditioning (Figures S6A and S6B) nor did we observe any physical withdraw symptoms during the test sessions. This suggests that in the current work, the MO-induced place preference, and changes in hippocampal place cell coding, reflect the rewarding rather than the withdraw effects of MO. In this case, the distinct effects of MA and MO may reflect differential cellular or molecular mechanisms (e.g. dopamine transporters versus opioid receptors), rather than a difference in reward versus aversive learning. However, the possibility remains that MO-conditioned mice experienced negative emotional withdrawal effects, as previous works suggest a single, high dose of an opioid can induce long-lasting opioid tolerance and increased pain sensitivity in naïve rodents (*91, 92*). Thus, it will be important for future work to consider drug-induced changes to hippocampal coding under both rewarding and aversive conditions.

## Acknowledgements

This work was supported by funding from NIDA DA042012, Office of Naval Research N000141812690, Simons Collaboration on the Global Brain 542987SPI, the Vallee Foundation and the James S McDonnell Foundation awarded to LMG. We thank A Diaz for animal husbandry assistance, B Heifets for CPP assistance and methamphetamine dosing, and X Xu for miniscope assistance.

## Author Contributions

YS and LMG conceptualized the experiments and analyses. YS collected and analyzed data. YS and LMG wrote the manuscript. LMG oversaw the project.

## Methods

### Subjects

All procedures were conducted according to the National Institutes of Health guidelines for animal care and use and approved by the Institutional Animal Care and Use Committee at Stanford University School of Medicine. For imaging experiments, Ai94;Camk2a-tTA;Camk2a- Cre (JAX id: 024115 and 005359) mice were used (n = 34 of mice in the study). We did not observe obvious epileptiform events in calcium activity or gross abnormal behavior in these mice (*93*). Male and female (Ctrl: 3 male and 4 female; MA: 6 male and 4 female; MO: 8 male and 4 female; MA+K: 2 male and 3 female) mice were group housed with same-sex littermates until the time of surgery. At the time of surgery, mice were 8 -12 weeks old (19 – 28 grams). After surgery mice were singly housed. Mice were kept on a 12-hour light/dark cycle and had *ad libitum* access to food and water in their home cages at all times. All experiments were carried out during the light phase.

### GRIN lens implantation and baseplate placement

Mice were anesthetized with continuous 1 – 1.5% isoflurane and head fixed in a rodent stereotax. A three-axis digitally controlled micromanipulator guided by a digital atlas was used to determine bregma and lambda coordinates. To implant the gradient refractive index (GRIN) lens above the CA1 regions of the hippocampus, a 1.8 mm-diameter circular craniotomy was made over the posterior cortex (centered at -2.30 mm anterior/posterior and +1.75 mm medial/lateral, relative to bregma). The dura was then gently removed and the cortex directly below the craniotomy aspirated using a 27- or 30-gauge blunt syringe needle attached to a vacuum pump under constant irrigation with sterile saline. The aspiration removed the corpus callosum above the hippocampal imaging window but left the alveus intact. Excessive bleeding was controlled using a hemostatic sponge that had been torn into small pieces and soaked in sterile saline. As determined using the Allen Brain Atlas (www.brain-map.org/), the unilateral cortical aspiration impacted part of the anteromedial visual area but the procedure left the primary visual area intact. The GRIN lens (0.25 pitch, 0.55 NA, 1.8 mm diameter and 4.31 mm in length, Edmund Optics) was then slowly lowered with a stereotaxic arm to CA1 to a depth of -1.53 mm relative to the measurement of the skull surface at bregma. A skull screw was placed on the contralateral side of the skull surface. Both the GRIN lens and skull screw were then fixed with cyanoacrylate and dental cement. Kwik-Sil (World Precision Instruments) was used to cover the lens at the end of surgery. Two weeks after the implantation of the GRIN lens, a small aluminum baseplate was cemented to the animal’s head on top of the existing dental cement. Specifically, Kwik-Sil was removed to expose the GRIN lens. A miniscope was then fitted into the baseplate and locked in position so that GCaMP6s expressing neurons and visible landmarks, such as blood vessels were in focus in the field of view. After the installation of the baseplate, the imaging window was fixed for the long-term in respect to the miniscope used during installation. Thus, for all imaging experiments, each mouse had a dedicated miniscope. When not imaging, a plastic cap was placed in the baseplate to protect the GRIN lens from dust and dirt.

### Methamphetamine and morphine conditioned place preference (CPP)

After mice had fully recovered from the baseplate surgery, they were handled and allowed to habituate to wearing the head-mounted miniscope by freely exploring an open arena for 20 minutes every day for one week. If the animal still showed muscle weakness (as indicated by the difficulty in holding their heads up towards the end of each session) at the end of the first week, they underwent an extra week of habituation. In parallel, animals were also habituated to mock intraperitoneal injections (needle poking) once a day for four days.

Conditioned place preference (CPP) sessions took place in a different room from the habituation sessions described in the previous paragraph. This dedicated CPP room contained salient distal visual cues, which were kept constant over the course of the entire experiment. The CPP apparatus consisted of two 25 x 25 cm compartments with distinct colors and visual cues (Figure 1A). The two compartments could be connected by a sliding door in the middle. The door opening was 6.5 cm wide, so that the mouse could easily run between the two compartments during miniscope recordings. The floors of the CPP compartments were covered with ∼500 ml of bedding to facilitate exploration. Mice were first habituated for two days (20 min/day) to the CPP apparatus with the door between the CPP compartments open and the miniscope mounted. Each mouse had its own dedicated miniscope for the entire duration of the CPP experiment, which ensured stable longitudinal recordings and facilitated image alignment across different sessions. Before any experiments started, the image quality for each animal was verified by adjusting the power of the excitation light and the focal plane. These scope parameters then remained fixed on the dedicated miniscope over the course of the entire CPP experiment.

After the two-day habituation, the pre-baseline session occurred on day 1, in which mice could run freely between the two connected CPP compartments for 20 minutes (with imaging). On day 2, the baseline session, the same experiment was repeated and the behavioral data used to assess the animals’ naturally preferred side (i.e. the compartment where the animal spent more time). Prior to the experiments, we defined an exclusion threshold, in which we would exclude any mouse that spent more than 75 % of their total time in one compartment in the baseline session; however, no mice reached this threshold. The subsequent conditioning sessions (3 sets, 6 days of pairings) were performed by confining the animal in one of the CPP compartments for 45 minutes immediately after a saline or drug administration. During the 45 minutes, we only imaged from the 15^th^ - 30^th^ minute in each mouse. For each set of conditioning sessions, saline was paired on the preferred side on the first conditioning day and the drug paired (MA at 2 mg/kg or MO at 20 mg/kg, injected intraperitoneally) on the non-preferred side on the second conditioning day. This conditioning process was repeated 3 times in total such that each animal received 3 saline pairings and 3 drug pairings. Similarly, in control (Ctrl) mice, saline was paired on both CPP compartments, starting with the preferred side. Note that for MA and MO mice, the naturally preferred side was equivalent to the saline-paired side and the non-preferred side was equivalent to the drug-paired side. 24 hours after the last conditioning session, animals were put back into the CPP environment with the two compartments connected again for 20 minutes with imaging to assess their post-conditioning preferences (defined as test 1). To assess whether any drug-induced preference was long-lasting, we performed a 2^nd^ post-conditioning test (defined as test 2) five days after test 1. The behavioral CPP score was defined as the time that the animal spent in the naturally non-preferred/drug side of the CPP apparatus in a test session minus the time it spent on the same side in the baseline session.

### Methamphetamine conditioned place preference (CPP) with ketamine blockade

A separate cohort of C57BL/6 mice (n = 8; 4 male and 4 female) were used to investigate whether ketamine blocks MA-induced place preference. For these experiments, we followed the same CPP protocol as MA-conditioning (described above), but ketamine (25 mg/kg) was administered intraperitoneally together with saline or MA in all of the conditioning sessions. For calcium imaging during MA CPP with ketamine, we used a separate cohort of Ai94;Camk2a- tTA;Camk2a-Cre mice (n = 5; 2 male and 3 female) and followed the same CPP procedure.

### Histology

Mice were transcardially perfused with 5 ml of phosphate buffered saline (PBS), followed by 25 ml of 4% paraformaldehyde-containing phosphate buffer. The brain was removed and left in 4% paraformaldehyde overnight. The next day, samples were transferred to 30% sucrose in PBS. At least 24 hours later, the brain was sectioned coronally into 40-µm-thick samples using a cryostat. Sections were mounted and cover-slipped with antifade mounting media with DAPI (Vectashield). Brain slice images were acquired using a ZEISS Axio Imager 2 fluorescence microscope under 10X or 20X magnification for both DAPI and GFP channels.

### Miniscope imaging data acquisition and initial batch processing

Technical details for the custom-constructed miniscopes and general processing analyses are described in (*48, 50*) and at miniscope.org. Briefly, this head-mounted scope had a mass of about 3 grams and a single, flexible coaxial cable to carried power, control signals, and imaging data to custom open source Data Acquisition (DAQ) hardware and software. In our experiments, we used Miniscope V3, which had a 700 μm x 450 μm field of view with a resolution of 752 pixels x 480 pixels (∼1 μm per pixel). Acquired data was packaged by the electronics to comply with the USB video class (UVC) protocol. The data was then transmitted via a Super Speed USB to a PC running custom DAQ software. The DAQ software was written in C++ and used Open Computer Vision (OpenCV) libraries for image acquisition. Images were acquired at ∼30 frames per second (fps) and recorded to uncompressed .avi files. The DAQ software also recorded the simultaneous behavior of the mouse through a high definition webcam (Logitech) at ∼30 fps, with time stamps applied to both video streams for offline alignment.

Miniscope videos of individual sessions were first concatenated and down-sampled by a factor of 2 using custom MATLAB scripts, then motion corrected using the NoRMCorre MATLAB package (*94*). To align miniscope videos across different sessions for the entire CPP experiment, we applied an automatic 2D image registration method (github.com/fordanic/image-registration) with rigid x-y translations according to the maximum intensity projection images for each session. The registered videos for each animal were then concatenated together in chronological order to generate a combined data set for extracting calcium activity. To extract the calcium activity from the large combined data set (> 10 GB), we used the Sherlock HPC cluster hosted by Stanford University to process the data across 8 – 12 cores and 600 – 700 GB of RAM. While processing this combined data set required significant computing resources, it enhanced our ability to track cells across sessions from different days (Figure 1). This process made it unnecessary to perform individual footprint alignment or cell registration across sessions.

To extract an individual neuron’s calcium activity, we adopted a newly developed method of extended constrained non-negative matrix factorization for endoscopic data (CNMF-E) (*53*). CNMF-E is based on the CNMF framework (*95*), which enables simultaneous denoising, deconvolving and demixing of calcium imaging data. A key feature includes modeling the large, rapidly fluctuating background, allowing good separation of single-neuron signals from background, and separation of partially overlapping neurons by taking a neuron’s spatial and temporal information into account (see (*53*) for details). After iteratively solving a constrained matrix factorization problem, CNMF-E extracted the spatial footprints of neurons and their associated temporal calcium activity. Specifically, the first step of estimating a given neuron’s temporal activity (a scaled version of dF/F, an oft used metric in calcium imaging studies) was to compute the weighted average of fluorescence intensities after subtracting the temporal activity of other neurons in the given neuron’s region of interest. A deconvolution algorithm called OASIS (*55*) was then applied to obtain the denoised neural activity and deconvolved spiking activity, as illustrated in Figure S1. These extracted calcium signals for the combined data set were then split back into each session according to their individual frame numbers.

The position and speed of the animal was determined by applying a custom MATLAB script to the animal’s behavioral tracking video. Time points at which the speed of the animal was lower than 2 cm/s were identified and excluded them from further analysis. We then used linear interpolation to temporally align the position data to the calcium imaging data.

To further validate the quality of cross-session alignment using the image registration, we analyzed the maximum projected images from each session after the registration with the colocalization analysis available via NIH ImageJ and the JACoP plugin (*96*) (Figure S1). In brief, the two images were background subtracted and a threshold automatically applied based on the Coste’s approach. A cytofluorogram was then plotted for each pixel pair based on their intensity. A Pearson’s coefficient can then be derived by calculating the best fitting regression line on the cytofluorogram. To test the statistical significance of the co-localization analysis, we used the Coste’s approach to randomly shuffle pixel blocks for one of the images 1000 times and obtained a distribution of concomitantly calculated shuffled coefficients. The 95^th^ percentile of the shuffled distribution were then determined as the significant threshold.

### Position-matching for comparisons of cell activity across sessions and CPP compartments

Analyses that compared hippocampal neuronal activity across different sessions (longitudinal comparisons) or across the two CPP compartments within the same session (transverse comparisons) could be influenced by biases in the animal’s occupancy, particularly due to the CPP related shift in spatial preference. To circumvent the effect of differences in occupancy on our analyses, we implemented a position-matched down-sampling protocol (*89*) when performing longitudinal or transverse comparisons of place cell activity. For down-sampling, we first binned the spatial arena into 1.8 x 1.8 cm non-overlapping bins. We then computed the number of position samples (frames) observed in each spatial bin and matched the number of samples between the two sessions (or compartments) by randomly removing position samples, and the corresponding neural activity, from the session (or compartment) with greater occupancy. Due to the stochastic nature of the down-sampling process, we repeated this procedure 50 times (unless otherwise specified) for each cell, and the final value for each cell was calculated as the average of all 50 iterations. This final value was then used to obtain the reported means or perform statistic comparisons. This protocol was applied to our analyses for all the within subject comparisons (both longitudinal and transvers). Specific details for each analysis are described in the corresponding methods and figure legends.

### Place cell analyses

#### Calculation of spatial rate maps

After we obtained the deconvolved spiking activity of neurons, we extracted and binarized the effective neuronal calcium events from the deconvolved spiking activity by applying a threshold (3 x standard deviation of all the deconvolved spiking activity for each neuron). The position data was sorted into 1.8 x 1.8 cm non-overlapping spatial bins. The spatial rate map for each neuron was constructed by dividing the total number of calcium events by the animal’s total occupancy in a given spatial bin. The rate maps were smoothed using a 2D convolution with a Gaussian filter that had a standard deviation of 2.

#### Spatial information and identification of place cells

To quantify the information content of a given neuron’s activity, we calculated spatial information scores in bits/spike (each calcium event is treated as a spike here) for each neuron according to the following formula (*97*),

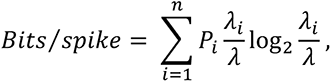

where *P*_*i*_ is the probability of the mouse occupying the i-th bin for the neuron, *λ*_*i*_ is the neuron’s unsmoothed event rate in the i-th bin, while *λ* is the mean rate of the neuron across the entire session. Bins with total occupancy time of less than 0.1 second were excluded from the calculation. To identify place cells, the timing of calcium events for each neuron was circularly shuffled 1000 times and spatial information (bits/spike) recalculated for each shuffle. This generated a distribution of shuffled information scores for each individual neuron. The value at the 95^th^ % of each shuffled distribution was used as the threshold for classifying a given neuron as a place cell, and we excluded cells with an overall mean calcium event rate less than 0.1 Hz. This threshold was roughly equal to the 5^th^ % of the mean event rate distribution for all neurons.

To avoid the potential influence of drug-induced change in occupancy on identifying place cells, we classified place cells by applying the position-matching protocol (described in the previous section) in addition to the above shuffling procedure. Namely, the full shuffling procedure was repeated for 20 times, each time on down-sampled neural activity obtained to match the position occupancy of the animal across the two CPP compartments. From these 20 repetitions, we calculated a place cell agreement index (PCI) for each neuron as

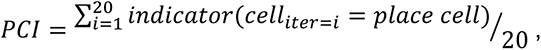

where indicator() is a function that returns 1 when the equation inside the parentheses holds, and 0 otherwise; i is the number of iterations (iter). We chose PCI ≥ 0.3 as the threshold for classifying a neuron as a place cell, based on both the visual inspection of place fields and the resulting proportion of place cells (∼70% of total neurons, union of the place cells from both CPP compartments), which was similar to conventional tetrode recordings from CA1 (*98, 99*). Note that compared with the regular shuffling method that does not perform position matching, the current approach results in slightly fewer cells classified as place cells. Although the place cells classified and used in the current study were all obtained through this position matching method, the general findings regarding the number of place cells described in the results (Figures 2B, 2C, and 2G; Figures 5D and 5E; Figures 6A and 6B) are robust and remain the same even without position matching.

#### Calculating the overlap between place cell populations

To measure the overlap and assess the distance between two place cell populations within each animal across different days, we calculated the Jaccard similarity index (*100*), defined as J(A, B) = ^*A* ∩ *B*^⁄_*A* ∪ *B*_ , where A and B are vectors containing numeric labels for cells that are place cells. In our analysis, A and B define two place cell populations from different sessions in a CPP compartment for a given mouse. By definition, the value of the Jaccard similarity index ranges from 0 to 1. Intuitively, if the two place cell populations are identical, the numerator will be equal to the denominator in the equation and thus gives a Jaccard index equals to 1.

#### Spatial and population vector (PV) correlations

Before calculating the spatial or PV correlation we first applied the position-matching protocol. The spatial correlation between two rate maps was calculated as Pearson’s correlation coefficient. For calculating population vector correlations, z-scored 2D rate maps of the selected group of neurons were first concatenated along the z dimension. A population vector (PV) was then extracted for each spatial bin (1.8 x 1.8 cm) along the z dimension. The PV correlation was then calculated as the Pearson’s correlation coefficient between the two PVs at the corresponding spatial bin across the two CPP compartments.

#### Quantification of place field size

We only measured place fields in neurons classified as place cells in a given environment. To measure the size of a given place field, the rate map was first binarized by applying a threshold of 50% of the peak event rate for the rate map. The place fields were then identified and extracted as connected objects from the binarized rate map. For place cells with more than one place field, we used the largest place field as the measurement.

### Reconstructing the mouse’s position using a naïve Bayes classifier

We used a naïve Bayes classifier to estimate the probability of animal’s location given the activity of all the recorded neurons. Speed filtered (> 2 cm/s), thresholded (3 x standard deviation of all the deconvolved spiking activity for each neuron), and binarized deconvolved spike activity (neuron.S in CNMF-E) from all neurons were first binned into non-overlapping time bins of 0.8 seconds. This time bin width was selected based on the overall decoding performance among all the bin width tested ranging from 0.2 to 1.6 seconds. The M x N spike data matrix, where M is the number of time bins and N is the number of neurons, was then used to train the decoder with an M x 1 vectorized location labels. The posterior probability of observing the animal’s position Y given neural activity X can then be inferred from the Bayes rule as:

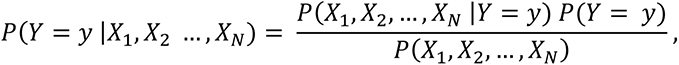

where X = (X_1_, X_2_, … X_N_) is the activity of all neurons, y is one of the spatial bins that the animal visited at a given time, and P(Y = y) is the prior probability of the animal being in spatial bin y. We used an empirical prior as it showed slightly better performance than a flat prior. P(X_1_, X_2_, …, X_N_) is the overall firing probability for all neurons, which can be considered as a constant and does not need to be estimated directly. Thus, the relationship can be simplified to

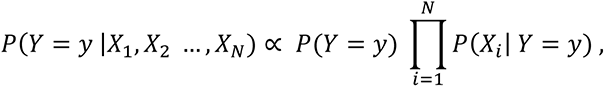

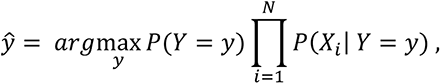

where *ŷ* is the animal’s predicted location, based on which spatial bin has the maximum probability across all the spatial bins for a given time. To estimate P(X_i_ | Y = y), we applied the built-in function of MATLAB fitcnb() to fit a multinomial distribution using the bag-of-tokens model with Laplace smoothing, which gave an estimation of distribution parameter for each neuron X_i_ at the given spatial bin y as:

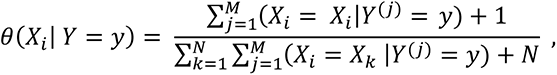

where, regardless of the Laplace smoothing, the numerator is the total number of spikes of neuron X_i_ for the time bins that the animal is at location y, and the denominator is the total number of spikes of all the neurons for the time bins that the animals is at location y. In addition, the above equation was weighted such that the normalized weights within a location bin sum to the prior probability for that location bin.

For the decoding analyses, we tested both the multinomial naïve Bayes classifier and a Poisson naïve Bayes classifier (*101*), the latter of which is often used in neural decoding. We found the multinomial based model outperformed the Poisson based model and provided higher decoding accuracy and a shorter time bin width associated with the best decoding performance (data not shown). As we are investigating the function of a small population of neurons, high decoding accuracy is important and thus, we chose to use the multinomial naïve Bayes classifier for our decoding analyses.

In addition, to reduce occasional erratic jumps in position estimates, we implemented a 2- step Bayesian method by introducing a continuity constraint (*101*), which incorporated information regarding the decoded position in the previous time step and the animal’s running speed to calculate the probability of the current location y. The continuity constraint for all the spatial bins Y at time t followed a 2D gaussian distribution centered at position y_t-1_, which can be written as:

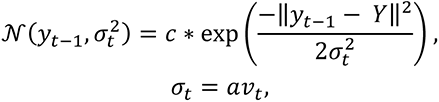

Where c is a scaling factor and v_t_ is the instantaneous speed of the animal between time t-1 and t. v_t_ is scaled by *a*, which is empirically selected as 2.5. The final reconstructed position with 2- step Bayesian method can be further written as:

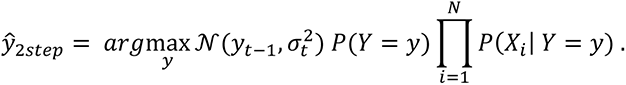

Decoded vectorized positions were then mapped back onto 2D space. The decoding error was calculated as the mean Euclidean distance between the decoded position and the animal’s true position, across all time bins. The overall improvement gained by implementing the 2-step Bayesian method was only ∼5-10%, which indicated that the method did not introduce any direct information regarding the animal’s current location to the decoder.

To compare position decoding performance across mice, we randomly down-sampled the number of neurons such that they matched across all mice (n = 150). We performed this down- sampling 50 times and trained the decoder and generated position predictions for each iteration.

The final result for a given mouse was then calculated as the average decoding performance across the 50 iterations. To examine the decoder’s performance using shuffled data, we circularly shuffled the neural data 100 times, shifting the spike times by 5 - 95% of the total data length randomly (i.e., 60 – 1140 seconds for a 20 min recording). The final result was then calculated as the average decoding performance across the 100 shuffles. To compare position decoding performance between the two CPP compartments, data for both training and predicting were matched for occupancy 50 times. The final result for a given compartment was then calculated as the average decoding performance across the 50 iterations.

For the knock out (KO) decoding analyses, we replaced the neural activity of cells that were ‘knocked out’ with vectors of zeroes. This knock out procedure was only applied to the data we used for predicting position locations, not for training, as ablating neurons directly from the training data will result in the model learning to compensate for the missing information. In addition, for the KO decoding analyses, the training data set was subject to the position matching protocol. The final result for each mouse was then calculated as the averaged decoding performance across all of the position matching iterations. For the random KO condition, we randomly selected the same number of neurons as in the KO condition for each iteration. For conditions in which the training and prediction data were both from the baseline or test sessions, we used the session with a lower time bias between the two compartments as the training data, which allowed us to use the session with the maximum amount of training data after position matching between the two CPP compartments. As in the regular decoding analysis, the KO decoding analysis provided reconstructed/predicted positions for an animal based on the neural activity; the CPP time could then be reconstructed from this position result. For KO analyses in Figures 3K and 3L, the decoder was trained using the baseline data, and rtPCp (retained place cells on the preferred side) were randomly selected for KO to match the number of disPCp from the test sessions data before making predictions. Both training and prediction data were occupancy matched between the two compartments. We then calculated the final decoding error for the predicted position by averaging the decoding errors from all the position matching iterations for each compartment.

### Principal component analysis

To perform principal component analysis (PCA), we treated the denoised calcium trace (neuron.C in CNMF-E, Figure S1C, blue trace) as neural activity, which reflects the true variance in calcium signals for the neural population compared to deconvolved spikes. The denoised calcium traces were filtered with a noise level threshold of 2 x (neuron.C_raw – neuron.C) for each neuron. The final data matrix of nFrames x nNeurons was fed into the PCA. To compare population coding of neurons before and after drug conditioning, we first ran PCA with the baseline data; then projected the data from the test session onto the principal component axis (eigenvectors) of the baseline results. This projection allowed us to visualize the population activity pattern in the same PCA space across different sessions. For visualization purpose, we plotted the top two PCs.

### Temporal clustering with k-means

#### Identification of temporal clusters

To cluster neurons according to the temporal information of their calcium signals, we employed a consensus k-means clustering method and calculated a consensus matrix measuring how frequently two samples were clustered together in multiple clustering runs with randomly sub-sampled data. We used the same data format as that described in the principal component analysis to compute the consensus matrix using sub-sampled data from the baseline session for 100 iterations. For each iteration, the calcium data from all the neurons was randomly sub-sampled at 90% of the total frame length, and neurons were partitioned into 5 groups (see determining optimal K below) using k-means clustering by calculating the pair-wise Pearson’s correlation coefficient from the sub-sampled calcium data. In each k-means run, there were 10 repeated clustering (replicates) using new initial cluster centroid positions. A consensus matrix was then obtained upon the completion of all the iterations by calculating the frequency with which two neurons were grouped together. Neuron pairs that showed the same cluster assignment across the highest number of iterations had a high consensus index value. On the other hand, neuron pairs that rarely clustered together had a low consensus index value. The final cluster was then determined using a hierarchical clustering method with complete linkage on the consensus matrix.

#### Determining the optimal number of clusters (K)

To estimate the optimal K value, we chose to search from 3 to 9 clusters. As we estimated the optimal K using either the local minima or maxima from the measurements described below, we expanded our K value search to range from 2 to 10 clusters for calculating the optimal K value. To determine the optimal K, we examined the performance of the K-clustered consensus matrix by visualizing the heatmap reorganized by linkage (Figure S3A). The consensus matrix can also be visualized by plotting it as a cumulative distribution function (CDF), as shown in Figure S3B. In the case of perfect clustering, the value of the consensus matrix will be either 0 or 1. Thus, the corresponding CDF would follow a Bernoulli distribution and the curve would be flat for intermediate values. Based on this principle, we employed a metric called the Proportion of Ambiguous Clustering (PAC) to estimate the optimal K (*102*). PAC is defined as the fraction of sample pairs with a consensus index value between [0.1, 0.9]. A low PAC value indicates the CDF curve is flat in the middle, thus allowing inference of the optimal K by identifying the lowest PAC (Figure S3C). In addition to PAC, we also implemented a cophenetic correlation based measurement to infer the optimal K (*103*). This measurement computes the Pearson’s correlation between the distance of neuron pairs in the consensus matrix and the cophenetic distance of neuron pairs obtained from the dendrogram tree used to reorder the consensus matrix by linkage. Thus, the cophenetic correlation measures how faithfully the dendrogram tree represents the distance between neuron pairs. A cophenetic correlation equal to 1 indicates a perfect consensus matrix. We plotted the cophenetic correlation as a function of K, with a range from 2 to 10, and selected the local maxima of cophenetic correlation as the optimal K (Figure S3D). As shown in Figures S3E and S3F, measurements from both methods gave similar inferences, pointing to the optimal K = 5.

#### Template matching method for sorting temporally defined clusters according to their spatial firing patterns

To compare temporally defined clusters across different animals, we took advantage of the ensemble spatial firing patterns for each temporally defined cluster and sorted them in into the following five groups: SW, southwest; SE, southeast; NW, northwest; NE, northeast, and center. These directions denoted the location of the peak in the ensemble activity in respect to the CPP environment. For each cluster, the peak of the ensemble activity in the two CPP compartments are always at a similar location (Figure S4) except for the center cluster, which only has a single peak at the junction between the two compartments. To perform unbiased and automated sorting, we developed a template matching method and computed the Pearson’s correlation between the ensemble spatial firing maps for each cluster and standard gaussian templates for each direction (Figure S4). Each standard template contains two simulated Gaussian fields at homotopic positions across the two compartments (Figure S4A). The field akin to the CPP midline had a variance of 12.5 cm while the field away from the midline had a variance of 25 cm. To avoid aberrant matching performance, the center group, which can be unambiguously identified, was held out during this matching process. The final matching result gave a 4 x 4 correlation matrix (Figure S4C), which we used to sort the ensemble activity of each cluster by assigning it to the group with the highest correlation. In most animals, temporal clusters fell into one of these non-overlapping groups with an unambiguous ordering. Occasionally, group ordering with maximum summed correlation was used if there was ambiguity in the group assignment.

#### Analyses on spatially defined clusters

To cluster neurons based on their spatial firing pattern, we performed a similar consensus K-means clustering for all the neurons using their spatial firing maps. Spatial maps of all the neurons were first vectorized and concatenated together into a nSpatialBins x nNeurons matrix and subsampled at 90% of all the spatial bins for each iteration. The clusters were then determined using a hierarchical clustering method with complete linkage on the consensus matrix. To compare the performance of spatially and temporally defined clusters on separating spatial information, we plotted the spatial correlation matrix of all neurons ordered by each cluster and computed silhouette values for each neuron. The silhouette value measured the similarity of a given neuron to other neurons within each cluster, compared to neurons in other clusters. The silhouette value *Sil*_*i*_ for the i-th neuron is defined as:

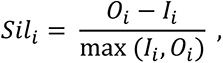

where *I*_*i*_ is the average within cluster distance of the i-th neuron, and *O*_*i*_ is the minimum average distance from the i-th neuron to neurons in other clusters, minimized over clusters. Silhouette value ranged from –1 to 1. A high silhouette value indicates that a given neuron is well matched to its own cluster, and poorly matched to other clusters. To determine which clustering method gave rise to a center cluster that better captured the neuron’s firing pattern during intercompartment traversals, we applied the template matching method to evaluate the fidelity of the spatial map for each center clustered neuron at the junction of the CPP environment. We then computed the Pearson’s correlation between the extracted portion from the center of each spatial rate map and a simulated Gaussian field with a variance of 2 (Figures S5F - S5H).

#### Measuring paired-wise anatomical distances

To measure the paired-wise anatomical distance for all disPCp in each mouse, we calculated the Euclidian distance between the centroid locations of each disPCp pair under the imaging window for each mouse. The centroid location of the neuron was obtained from the CNMF-E framework (neuron.centroid), denoting the center coordinates for each ROI contour. For each disPCp, we quantified an averaged intra- vs. inter-cluster distances based on the cluster assignment for all disPCp. The final result for each cluster was averaged across all disPCp that belonged to the same cluster. We expected that the inter-cluster distance would be larger than the intra-cluster distance, if functionally-defined disPCp clusters are anatomically clustered.

### Statistical Analysis

All the analyses and statistical tests were performed using MATLAB (2017b and 2020a). Data are presented as mean ± SEM or median ± interquartile range (IQR), as indicated. For statistical comparisons between groups, the data were checked for normality using Shapiro-Wilk tests. If the criteria for normality were met, an unpaired t-test was used to compare two groups; otherwise, a Wilcoxon rank-sum test was used. For statistical comparisons across more than two groups, One-Way ANOVA and related multiple comparison tests were used. For paired statistical comparisons, a paired t-test was used if the data followed a normal distribution; otherwise, a Wilcoxon sign-rank test was used. All tests were two-tailed unless otherwise specified. Most of the statistical tests were performed based on each animal or each session, as indicated in the appropriate figure legend. However, for paired observations across different sessions that involved large numbers of neurons from relatively small number of animals, we employed a linear mixed-effects model (LME) (‘fitlme’ in MATLAB) to perform hypothesis testing. The idea behind LME is to take the inherent independence in the data, such as neurons from the same mouse, into consideration when conducting statistical modeling and hypothesis testing. We fitted an LME by using different sessions (baseline vs. test) as a fixed effect and different mice as a random effect, so that we could account for the large sample size (number of neurons) while still controlling for the number of animals. In all experiments, the level of statistical significance was defined as *p* ≤ 0.05.

---

**Figure S1, related to Figure 1.**
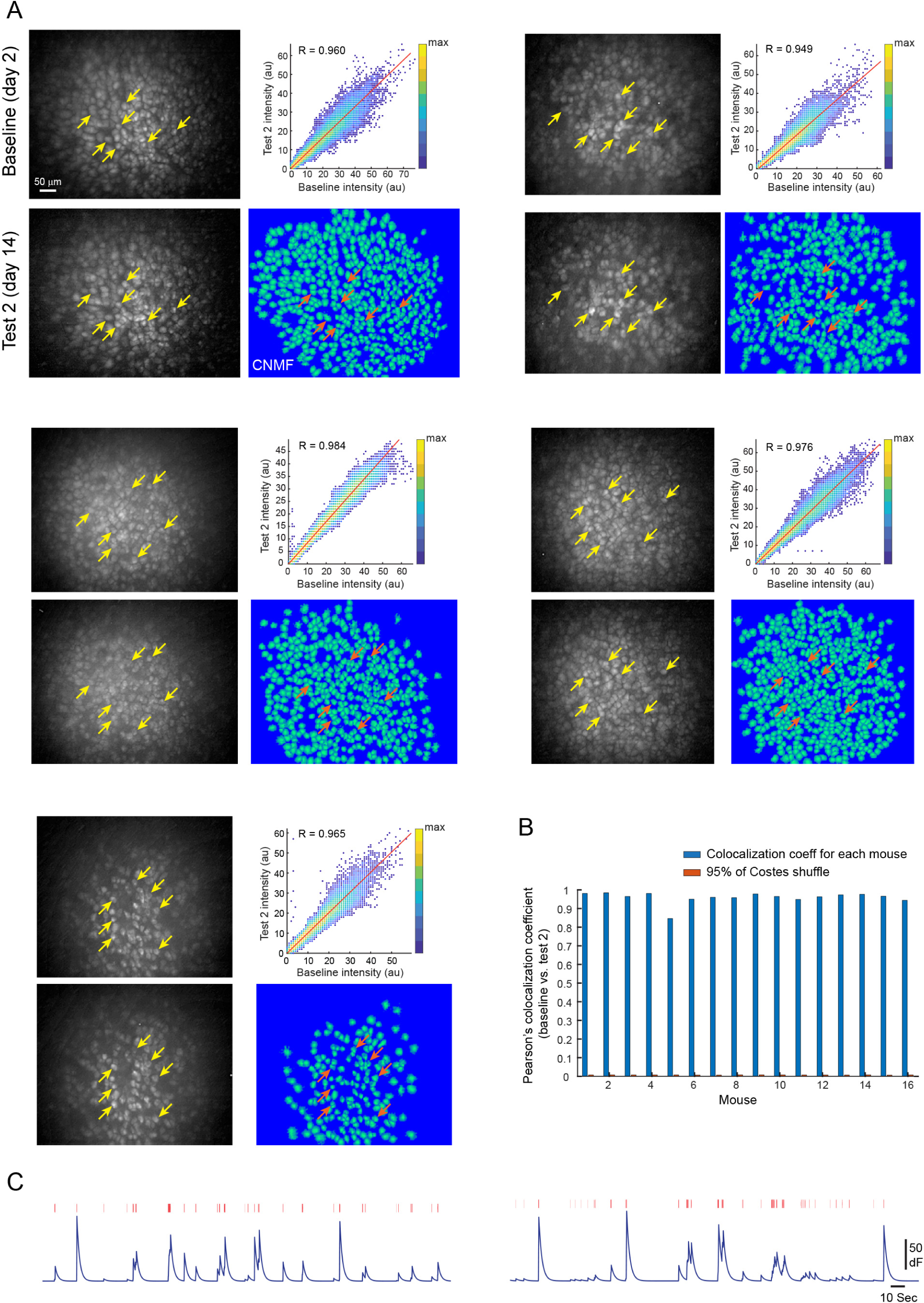
Long-term single cell tracking of hippocampal CA1 neurons using a miniscope. **A**, Examples of single cell tracking from five individual mice. Each panel block is a mouse. For each block, maximum intensity projected images of CA1 calcium imaging for baseline and test 2 sessions are shown on the left. Yellow arrows point to representative tracked neurons. Top right, cytofluorogram of colocalization analysis between baseline and test 2 images on the left. Fluorescence intensity for corresponding pixels of the two images are plotted. The figure is color coded for maximum (yellow) and minimum (blue) values with warmer colors indicating higher data point frequency. R value shows the Pearson’s colocalization coefficient between the two images. Bottom right, CNMF-E spatial footprints of neurons extracted from the aligned and concatenated data. Orange arrows point to the extracted footprints corresponding to the neurons indicated by yellow arrows on the left. **B**, Colocalization analyses between baseline and test 2 sessions for all Ctrl and MA mice (n = 16). Pearson’s colocalization coefficient (blue bar) and the 95% quantile of the Coste’s shuffled correlation coefficient (orange bar, see STAR methods) are shown side by side for each mouse. The averaged Pearson’s colocalization coefficient and the 95% quantile of the Coste’s shuffled correlation coefficient across all the mice are 0.96 ± 0.008 and 0.007 ± 0.0001 (mean ± SE), respectively. **C,** Representative denoised calcium signal traces (shown in dark blue) and deconvolved inferred spikes (shown in red bar) extracted from CNMF-E.

**Figure S2, related to Figure 2 and 3.**
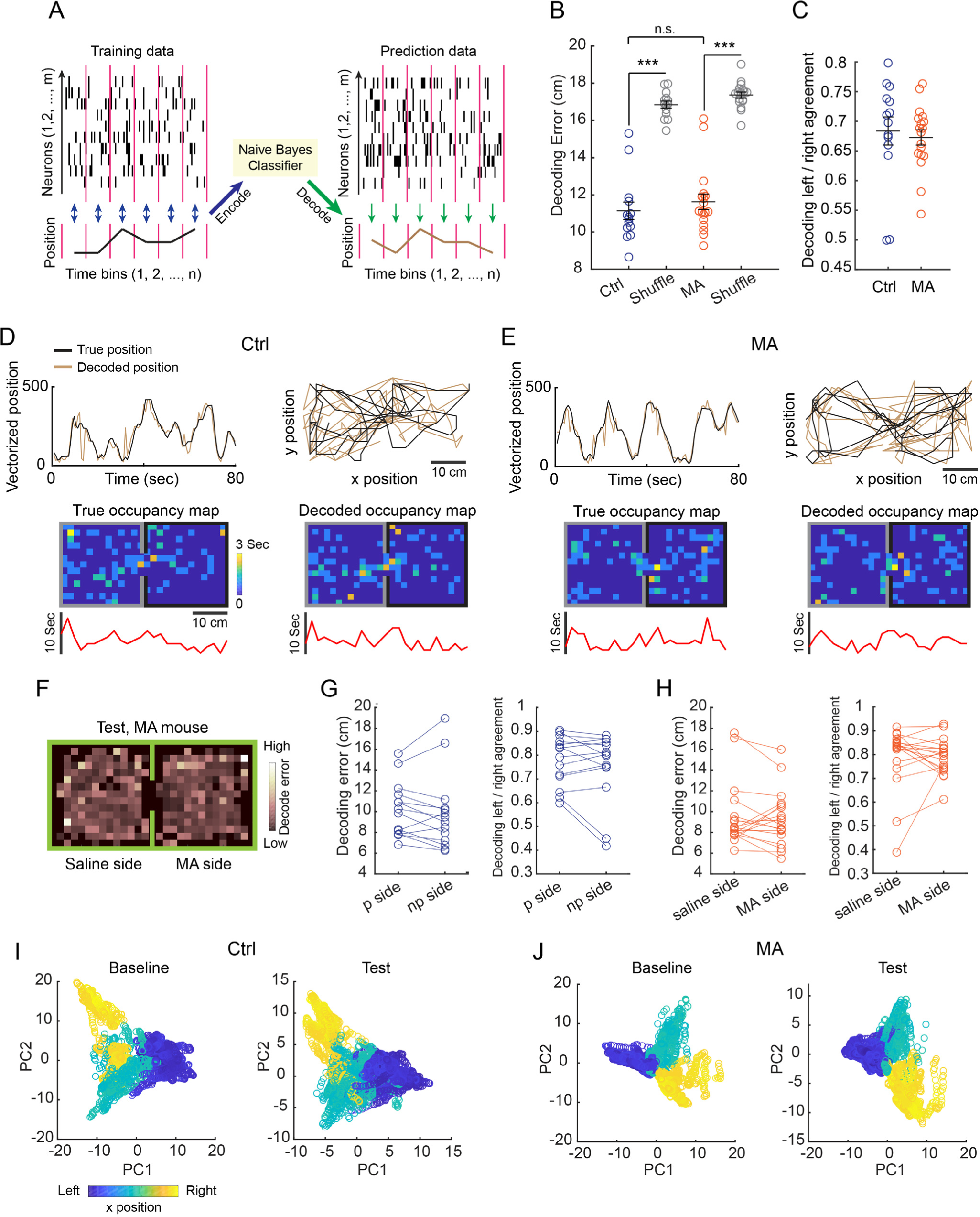
MA-induced place cell remapping does not impair spatial decoding performance. **A**, Schematic illustration of training (encoding) and prediction (decoding) processes for the naïve Bayes classifier. Neural and behavioral data were binned at 800 ms. The black trace indicates the true position of the animal, while the brown trace indicates the predicted/decoded position. **B - C**, Decoding performance for data from Ctrl mice, MA mice and the shuffled condition. The decoder was trained using baseline data in which the number of neurons was randomly down-sampled such that a comparable number was used across different animals. Predictions were then made using data from test sessions with the same down-sampling approach. We performed this down-sampling 50 times for each mouse and the final results were averaged from these 50 iterations. While decoding errors were significantly better than the shuffle for both groups (panel B, Ctrl vs. shuffle: 11.14 ± 0.47 vs. 16.85 ± 0.19 cm, mean ± SEM, p = 1.22 x 10^-4^; MA vs. shuffle: 11.63 ± 0.42 vs. 17.36 ± 0.16 cm, Z = -3.72, p = 1.96 x 10^-4^, two-tailed sign-rank test, n = 14 and 18 sessions. Note: two test sessions for each animal), there were no differences in either the decoding errors (panel B, Ctrl vs. MA: Z = -1.42, p = 0.15, two-tailed rank sum test) or decoding agreements (panel C, Ctrl 0.68 ± 0.02 vs. MA 0.67 ± 0.01, Z = 1.04, p = 0.30, two-tailed rank sum test) between Ctrl and MA mice. **D**, Top, an example of the true vs. decoded spatial position of a Ctrl mouse in 1D (left) and 2D (right) over an 80-second long period. Bottom, example occupancy map of the true vs. decoded spatial position for the same Ctrl mouse and data shown on top. The red trace below the occupancy map shows the summed time for each column of spatial bins in the occupancy map above. **E**, Organized as in (D), for a representative MA mouse. **F**, A 2D visualization of the decoding error in each spatial bin of the CPP environment from a test session of an MA mouse. Brighter colors indicate higher decoding errors. **G**, Comparisons of decoding error (preferred (p) side: 10.19 ± 0.71, mean ± SEM, non-preferred (np) side: 9.9 ± 0.99 cm, p = 0.50, two-tailed sign-rank test, n = 14) and decoding agreement (p side 0.78 ± 0.03, np side: 0.75 ± 0.04, p = 0.81, two-tailed sign-rank test, n = 14) across the two CPP compartments in test sessions of Ctrl mice. The decoder was trained using baseline data and predictions were made using test session data. Each animal’s occupancy was matched between the two compartments for both the training and prediction. We performed this occupancy matching 50 times for each mouse and the final results were averaged from these 50 iterations. As the comparisons were within subjects, the analyses were performed without down- sampling the number of neurons. **H**, Organized as in (G), comparisons of decoding error (mean ± SEM, saline side: 9.56 ± 0.73, MA side: 9.33 ± 0.64 cm, Z = 0.63, p = 0.53, two-tailed sign-rank test, n = 18) and decoding agreement (saline side: 0.79 ± 0.03, MA side: 0.79 ± 0.02, Z = 0.46, p = 0.65, two-tailed sign-rank test, n = 18) in test sessions of MA mice. **I – J**, Principal component analysis (PCA) revealed stable neural activity dynamics between the baseline and test sessions for both Ctrl (I) and MA (J) mice. An example mouse for each group was shown here. Data were first matched for occupancy between the baseline and test sessions. PCA was then performed using thresholded de-noised calcium signals from all neurons in the baseline session (a matrix with number of frames x number of neurons). To compare PCA results between the baseline and test sessions, data from test sessions were projected onto the principal axes of baseline PCs. To better visualize the structure of the data, we plotted data from the far left, far right, and middle of the CPP environment (10% of the total horizontal length for each). In both Ctrl and MA mice, the overall structure of PCA applied to the baseline and test sessions followed the same shape and orientation.

**Figure S3, related to Figure 4.**
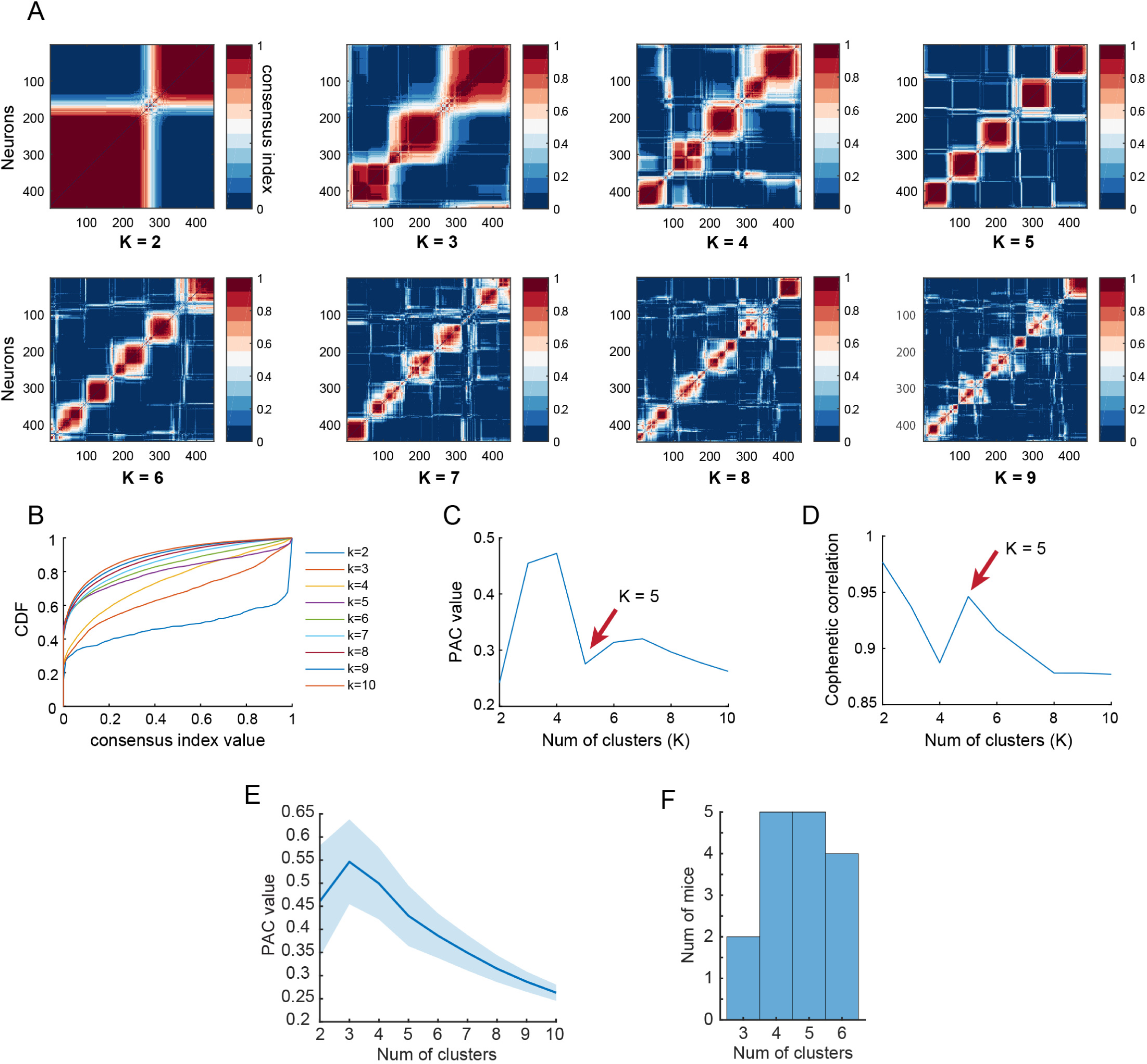
Determining the optimal number of temporal clusters using a consensus k-means method. **A**, Consensus matrices obtained at given numbers of clusters (K) using data from the baseline session of an MA mouse. The matrices are colored code for high (red) and low (blue) consensus index values. The consensus matrix was computed based on k-means clustering using randomly sub-sampled data with 100 iterations. Neuron pairs which showed the same cluster assignment in many of the iterations will have a high consensus index value. On the other hand, neuron pairs which rarely show the same cluster assignment will have a low consensus value. The matrices shown in the figure were re-organized using a hierarchical clustering method with complete linkage. **B**, An example cumulative distribution function (CDF) plot of the consensus index values, at different numbers of clusters, obtained from the same data set in (A). A consensus index value CDF with perfect clustering should follow a Bernoulli distribution and show a flattened curve in the middle. **C**, An example mouse of using the Proportion of Ambiguous Clustering (PAC) value to determine the optimal number of clusters on the baseline data. PAC is calculated by using the proportion of data points with a consensus index value between 0.1 – 0.9 on the CDF plot (see panel B). The optimal number of clusters is indicated by the local minima (red arrow). **D,** An example mouse of using the cophenetic correlation to determine the optimal number of clusters on the baseline data, which is indicated by the local maxima (red arrow). **E**, Averaged PAC value for the baseline data of Ctrl and MA mice (n = 16 mice) at given numbers of clusters. Note the elbow shape is at K = 5. Data are plotted with mean (dark blue line) ± SE (light blue shading). **F**, Histogram showing the number of mice (both Ctrl and MA) with their optimal number of clusters (n = 16 mice), as determined by the cophenetic correlation. Together, given the data plotted in (E) and (F), the optimal number of clusters K = 5 was used for all analyses.

**Figure S4, related to Figure 4.**
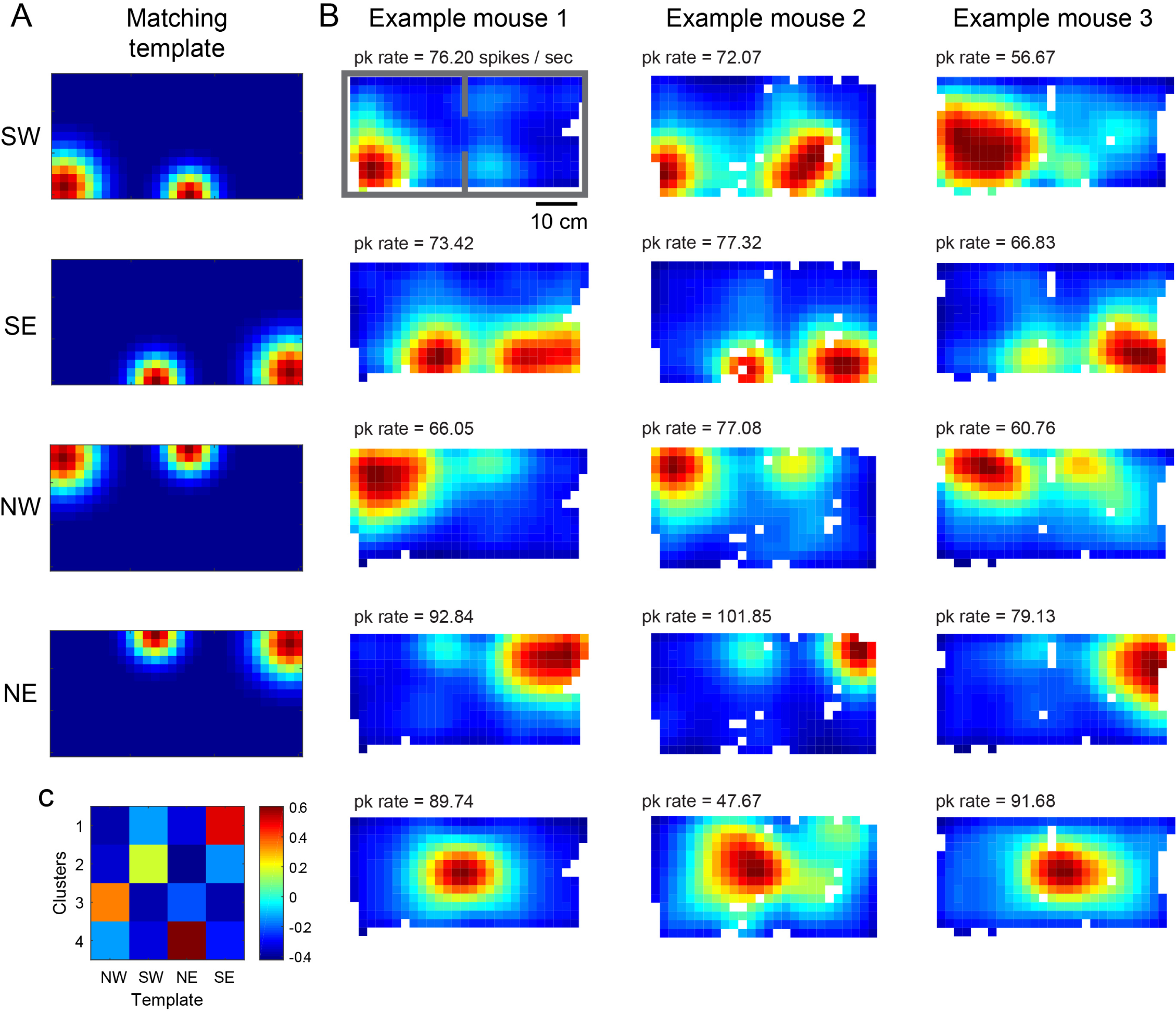
A template matching method to sort temporal clusters across animals using their spatial firing patterns. **A**, Gaussian spatial firing pattern templates used for matching (see STAR methods for details). SW, southwest; SE, southeast; NW, northwest; NE, northeast. Templates are color coded for minimum (blue) and maximum (red) values to mimic the spatial firing pattern of the ensemble activity observed in temporal clusters (shown in B). In brief, the matching process is to compute a Pearson’s correlation coefficient between an ensemble spatial rate map from a given temporal cluster (i.e., a summed rate map of all neurons from the cluster, shown in B) and the gaussian templates, then find the template with the highest correlation. To avoid aberrant matching performance, the ‘center’ group, which can be unambiguously identified, was held out during this matching process. **B**, Ensemble spatial rate maps for the five temporal clusters from three example mice. The rate maps are color coded for minimum (blue) and maximum (red) values with peak Ca^2+^ event rates indicated on the top left and organized according to the best matched template in (A). **C**, The correlation matrix resulting from the template matching process from an example mouse. Each cluster can be unambiguously assigned to a template based on the highest correlation value in each column.

**Figure S5, related to Figure 4.**
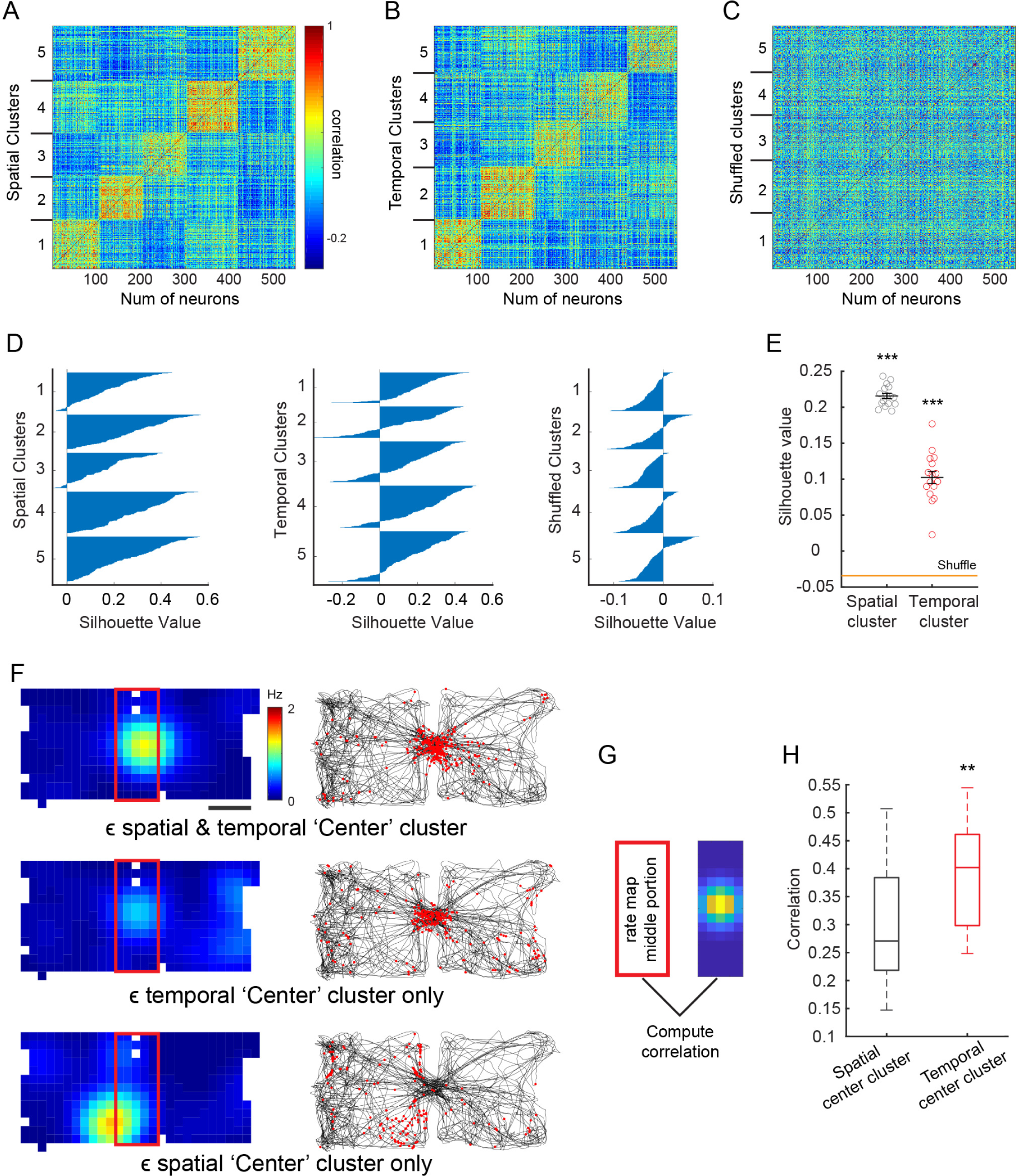
Temporally defined clusters are able to resolve spatial information and capture neural activity associated with inter-compartment traversals. **A**, A spatial correlation matrix sorted by spatially defined clusters. The matrix is computed from the baseline session of one representative mouse by calculating pair-wise spatial rate map correlations between neurons. The matrix is color coded for minimum (blue) and maximum (red) Pearson’s correlation values. Each column or line represents a neuron. Clusters are determined using a consensus k-means method. **B**, A spatial correlation matrix, as in (A), sorted by temporally defined clusters. Clear clustering is still visually observable in the matrix. **C**, A spatial correlation matrix sorted by a shuffled cluster order. **D,** Silhouette plots for (A), (B), and (C), respectively. The silhouette value is a measure of how similar a neuron is to its own cluster compared to other clusters; a high silhouette value indicates better clustering performance. The silhouette plot shows the data split across five clusters. Each line of a given cluster represents the silhouette value of an individual neuron. **E,** Silhouette values for spatially (grey, mean ± SEM: 0.22 ± 0.0037) and temporally (red, 0.10 ± 0.0087) defined clusters derived from the baseline spatial correlation matrix for each animal. Both spatially and temporally defined clusters showed significant higher silhouette values than the shuffled (orange line, -0.034 ± 0.0025) clusters (Spatial vs. shuffle: t(15) = 62.96, p = 1.34 x 10^-19^; Temporal vs. shuffle: t(15) = 18.79, 7.75 x 10^-12^, respectively, two-tailed paired t-test, n = 16 mice, Ctrl + MA). This indicates temporally defined CA1 neural clusters carry significant information regarding spatial activity. **F,** Example neurons in a representative mouse that belong to spatially or temporally defined ‘center’ clusters in the baseline session. The top example shows a neuron that belongs to both spatially and temporally defined ‘center’ clusters. The middle example shows a neuron that belongs only to the temporally defined ‘center’ cluster. The bottom example shows a neuron that belongs only to the spatially defined ‘center’ cluster. For each cell, the spatial firing rate map (left, color coded for minimum, blue, and maximum, red, values) and raster plot (right, red dots) on top of animals running trajectory (black traces) are shown. **G,** Illustration of the template matching method used to measure the correlation between the middle portion of a rate map (red box in F) and a gaussian template (right) for ‘center’ clustered neurons. **H**, Template matching analysis for spatially vs. temporally defined ‘center’ clusters. The Pearson’s correlation was computed for each ‘center’ clustered neuron in the baseline session for each animal. The temporal ‘center’ cluster (red, median: 0.40) showed significantly higher correlations with the gaussian template than the spatial ‘center’ cluster (black, median: 0.27) (z = -2.74, p = 0.0061, two-tailed sign-rank test, n = 16). Box plots show median, interquartile and full range of averaged values for each mouse. This result indicates that the temporal ‘center’ cluster was better at capturing neural activity corresponding to an animal’s inter-compartment traversals. This is likely due to the spatially defined clusters were grouped purely based on the location proximity of their firing fields rather than synchronized activity associated with a particular event.

**Figure S6, related to Figure 4 and 6.**
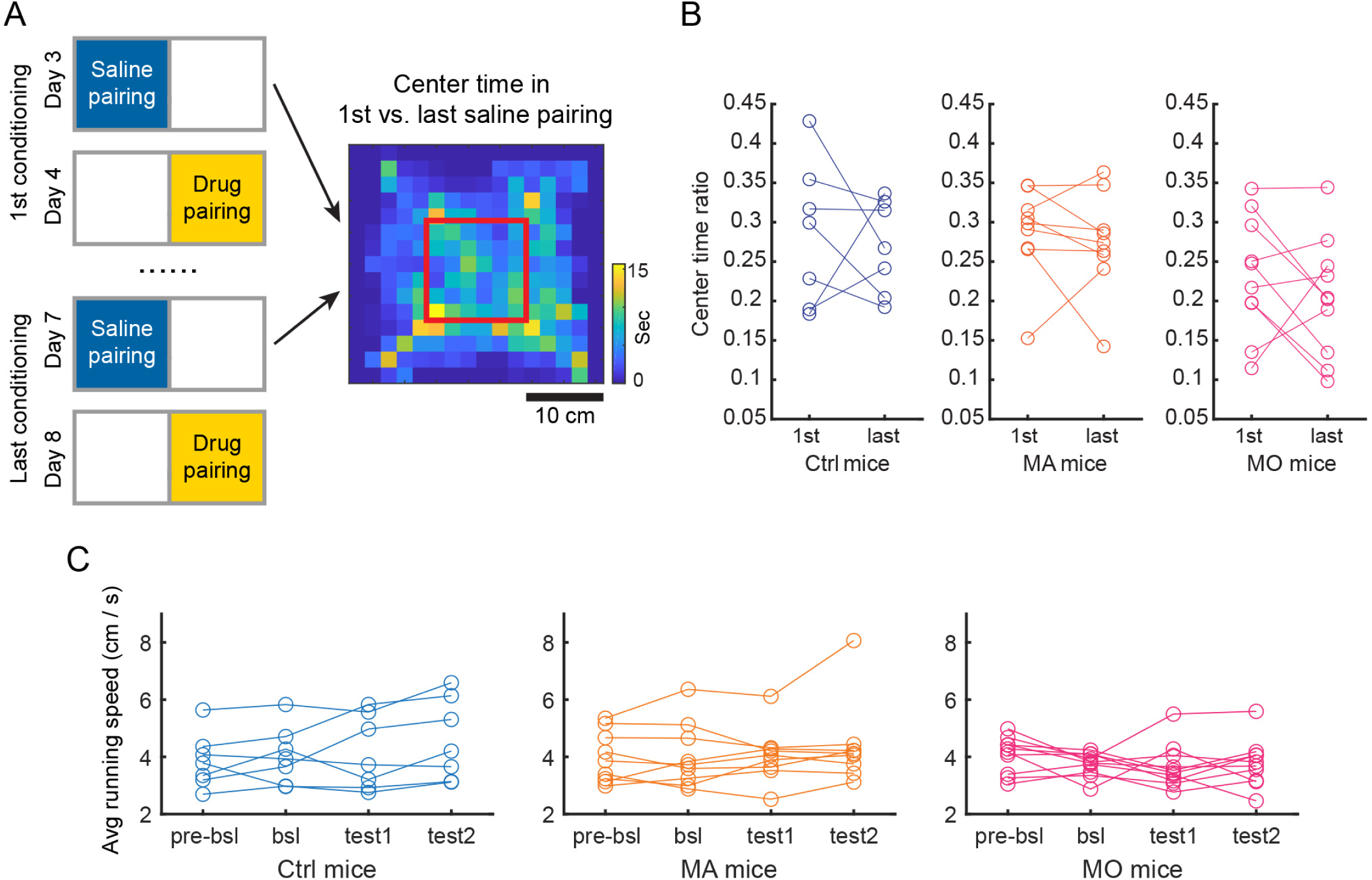
Repeated drug-conditioning did not change an animal’s occupancy time in the center of the arena nor the running speed. **A**, Left, schematic shows the timeline for the 1^st^ and last conditioning sessions. Note there was no drug experience prior to the 1^st^ saline pairing. Right, a representative example from an individual mouse of the occupancy time that the animal spent in the center of the arena, denoted by the red box. The occupancy map is color coded for minimum (blue) and maximum (yellow) values. To compare occupancy across drug conditioning, time spent in the center of the arena was calculated for both the 1^st^ and the last saline pairings, between which there were two drug-pairing sessions. **B**, Quantification of the proportion of time spent in the center of the arena in 1^st^ vs. last saline pairing (1^st^ vs. last, Ctrl: 0.29 ± 0.03 vs. 0.27 ± 0.02, mean ± SEM; MA: 0.29 ± 0.02 vs. 0.27 ± 0.02; MO: 0.23 ± 0.02 vs. 0.20 ± 0.02). There were no significant differences within the Ctrl, MA or MO group (Ctrl: t(6) = 0.44, p = 0.67; MA: t(8) = 0.67, p = 0.52; MO: t(9) = 1.05, p = 0.32, two-tailed paired t-test). This suggests that MA and MO drug-conditioning did not significantly change the stress or anxiety level of the mice, typical behavioral characteristics indicative of drug withdrawal. **C**, Drug conditioning did not change the running speed of mice in any group (Ctrl: F(3,24) = 0.47, p = 0.71; MA: F(3,32) = 0.22, p = 0.88; MO: F(3,36) = 0.65, p = 0.59; One-way ANOVA). Each line indicates data from an individual animal.

**Figure S7, related to Figure 6.**
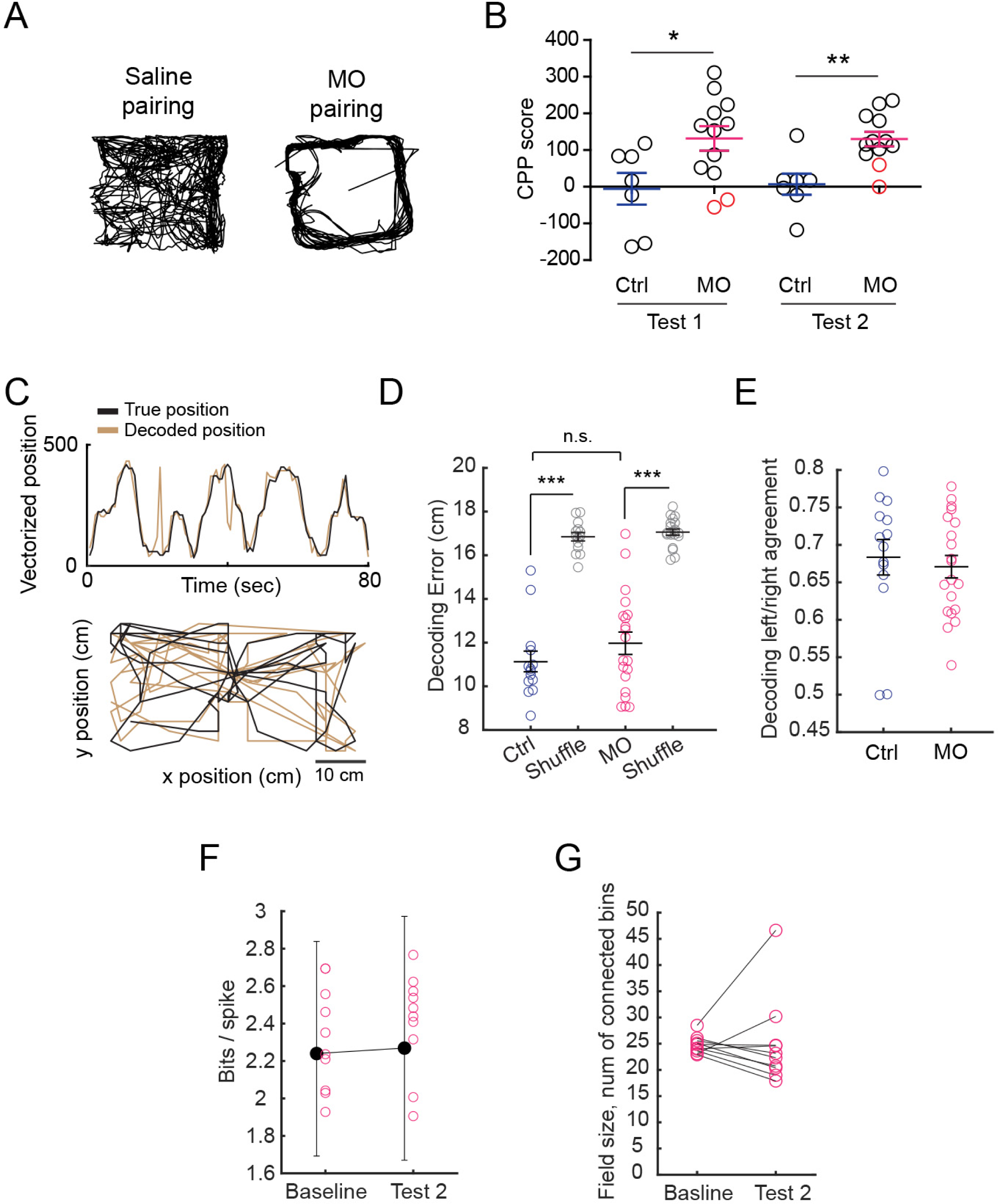
Morphine (MO) CPP behavior and spatial decoding accuracy. **A**, Similar to MA, administration of MO induced stereotypic circling behavior in mice, which was not observed after administration of saline. An example for one mouse is shown, with the black trace indicating mouse’s trajectory during a representation saline and MO pairing session. **B**, Behavioral CPP scores (time spent in the non-preferred/drug-paired side in the test compared to baseline session) for MO mice. MO mice showed a significant preference for the drug-paired side after conditioning compared with Ctrl mice for both test 1 (Ctrl vs. MA: -5.62 ± 43.28 vs. 131.80 ± 33.48 s, mean ± SEM; t(17) = 2.50, p = 0.023, two-tailed unpaired t-test. n = 7 and 12, respectively) and test 2 (Ctrl vs. MA: 6.89 ± 28.67 vs. 130.10 ± 19.82 s; t(17) = 3.64, p = 0.0020, two-tailed unpaired t-test). Red-colored data points indicate a neutral response to MO (defined as a negative CPP score). Data from these mice were excluded from further analyses. **C**, An example of the true vs. decoded spatial position of an MO mouse in 1D (top) and 2D (bottom) over an 80-second long period. **D-E**, Comparisons of decoding error (D) and decoding agreement (E) from Ctrl mice, MO mice and the shuffled condition. The decoder was trained using baseline data and predictions were made using test session data. For this analysis, the number of neurons was randomly down-sampled such that a comparable number was used across different animals. While decoding error was significantly lower than the shuffle condition in both Ctrl and MO mice (panel D, Ctrl vs. shuffle: 11.14 ± 0.47 vs. 16.85 ± 0.19 cm, mean ± SEM, p = 1.22 x 10^-4^; MO vs. shuffle: 11.97 ± 0.51 vs. 17.06 ± 0.14 cm, Z = -3.92, p = 8.86 x 10^-5^, two-tailed sign-rank test, n = 14 and 20 sessions. Note there were two test sessions for each animal), there were no differences in either the decoding error (panel D, Ctrl vs. MO: Z = -1.00, p = 0.32, two-tailed rank sum test) or decoding agreement (panel E, Ctrl 0.68 ± 0.02 vs. MO 0.67 ± 0.02, Z = 0.82, p = 0.41, two-tailed rank sum test) between Ctrl and MO mice. **F**, Compared with baseline, MO conditioning did not change the spatial information of disPCp place fields on the non-preferred/MO side in test 2 session (F(1,980) = 2.18, p = 0.14; linear mixed effect model, n = 491 cells from 10 mice). Black bar plots indicate the median and interquartile range of values for all the disPCp from all mice. Colored circles indicate the mean values for each animal. **G**, The field size of disPCp maintained the same before and after MO conditioning on the non-preferred/MO side (median field size bsl vs. test 2: 24.49 vs. 22.73 bins, p = 0.56, two-tailed sign-rank t-test, n = 10).

**Figure S8, related to Figure 6.**
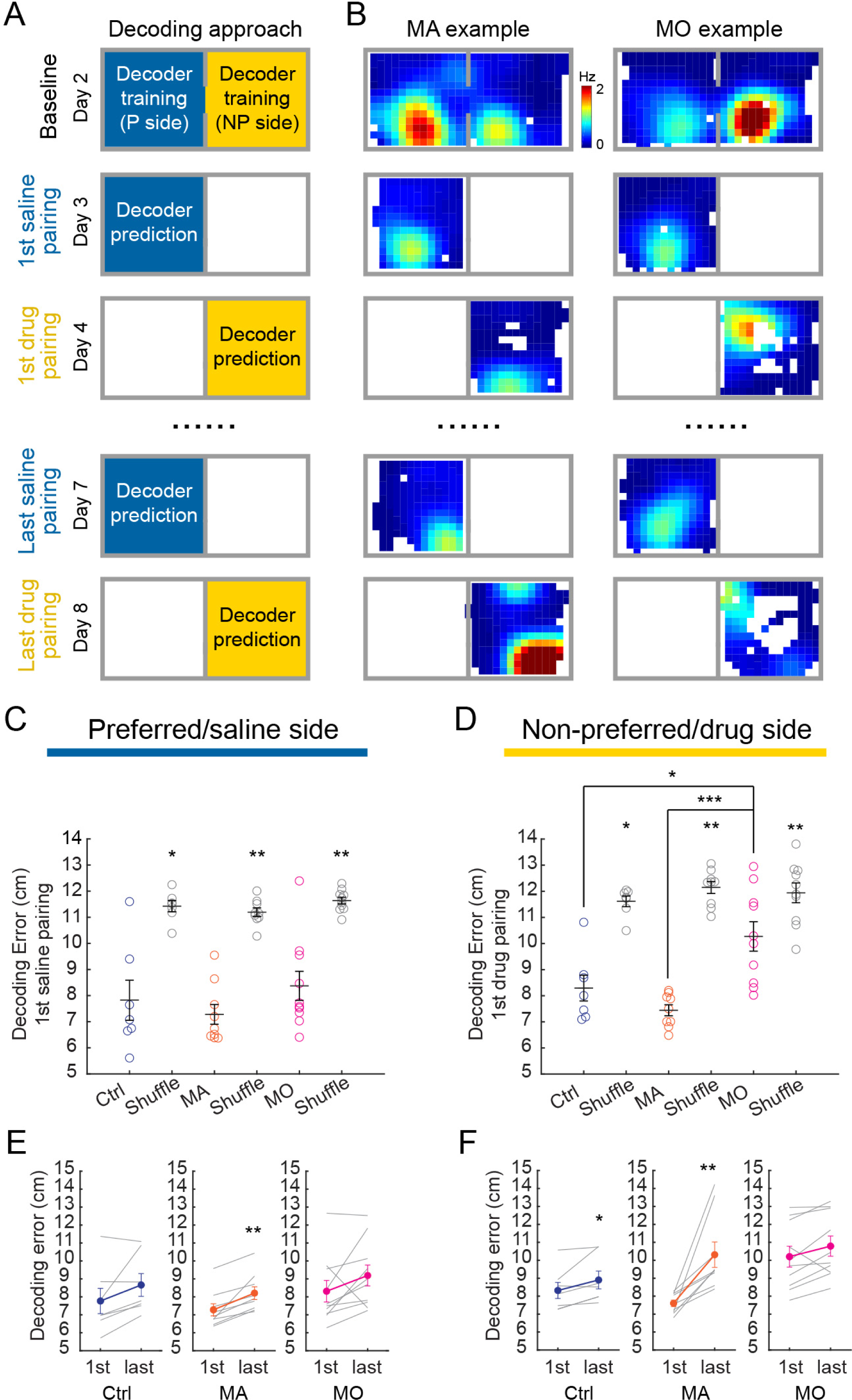
Effect of MA / MO administration on place cell representations in the conditioning sessions. **A**, Decoding approach applied to the conditioning sessions. Decoders were trained using either the preferred side data or the non-preferred side data from the baseline session. The decoders were then used to make position predictions for the corresponding conditioning sessions on the same side. Only the 1^st^ and the last (3^rd^) sets of conditioning were used to make the predictions and analytical comparisons. Note there was no drug experience prior to the 1^st^ saline pairing. **B**, Example place cells from MA and MO mice imaged across baseline and conditioning sessions. Each column is a cell. Spatial firing rate maps (color coded for minimum, blue, and maximum, red, values) for each neuron are shown. **C**, Comparisons of decoding errors for the 1^st^ saline pairing across groups. The decoder was trained using the preferred side data in baseline, with the number of neurons randomly down- sampled such that a comparable number was used across different animals. Although the decoding performance for each group was higher than the shuffle condition (Ctrl vs. shuffle: 7.82 ± 0.77 vs. 11.42 ± 0.21 cm, mean ± SEM, p = 0.031; MA vs. shuffle: 7.28 ± 0.38 vs. 11.19 ± 0.16 cm, p = 0.0039; MO vs. shuffle: 8.37 ± 0.56 vs. 11.63 ± 0.13 cm, p = 0.0039, two-tailed sign-rank test, n = 7, 9, and 10, respectively), decoding error was not different between groups (Ctrl vs. MA: p = 0.61; Ctrl vs. MO: p = 0.36, two-tailed rank sum test). **D**, Comparisons of decoding errors for the 1^st^ drug pairing across groups. Although the decoding performance for each group was higher than the shuffle condition (Ctrl vs. shuffle: 8.29 ± 0.49 vs. 11.62 ± 0.21 cm, mean ± SEM, p = 0.016; MA vs. shuffle: 7.44 ± 0.21 vs. 12.15 ± 0.23 cm, p = 0.0039; MO vs. shuffle: 10.27 ± 0.56 vs. 11.94 ± 0.38 cm, p = 0.0020, two-tailed sign-rank test, n = 7, 9, and 10, respectively), decoding errors in MO mice were significantly larger than Ctrl and MA mice (Ctrl vs. MO: p = 0.025, MA vs. MO p = 8.66 x 10^-5^, two-tailed rank sum test). This suggests that place cells showed more significant remapping in MO mice, compared to MA mice, during the 1^st^ drug pairing. **E**, Comparison of decoding error between the 1^st^ and last saline-pairing within each group. The decoder was trained using the preferred side data in baseline and predictions were made using the occupancy matched data between the 1^st^ and last saline pairing. There was a significant increase in the decoding error in MA mice but not in Ctrl or MO mice (Ctrl: 7.77 ± 0.70 vs. 8.66 ± 0.63 cm, mean ± SEM, p = 0.078; MA: 7.28 ± 0.34 vs. 8.21 ± 0.35 cm, p = 0.0039, MO: 8.31 ± 0.60 vs. 9.19 ± 0.58 cm, p = 0.064, two-tailed sign-rank test, n = 7, 9, and 10 mice). Colored dots and lines correspond to mean ± SEM, with data from individual mice shown as grey lines. **F**, Comparison of decoding error between the 1^st^ and last drug-pairing for each group. There was a significant increase in the decoding error in both the Ctrl and MA mice, but not in the MO mice (Ctrl: 8.32 ± 0.45 vs. 8.90 ± 0.50 cm, p = 0.031; MA: 7.60 ± 0.19 vs. 10.31 ± 0.71 cm, p = 0.0039, MO: 10.20 ± 0.58 vs. 10.79 ± 0.56 cm, p = 0.064, two-tailed sign-rank test, n = 7, 9, and 10 mice).

